# Dopaminergic medication reduces striatal sensitivity to negative outcomes in Parkinson’s disease

**DOI:** 10.1101/445528

**Authors:** Brónagh McCoy, Sara Jahfari, Gwenda Engels, Tomas Knapen, Jan Theeuwes

## Abstract

Reduced levels of dopamine in Parkinson’s disease (PD) contribute to changes in learning, resulting from the loss of midbrain dopamine neurons that transmit a teaching signal to the striatum. Dopamine medication used by PD patients has previously been linked to either behavioral changes during learning itself or adjustments in approach and avoidance behavior after learning. To date, however, very little is known about the specific relationship between dopaminergic medication-driven differences during learning and subsequent changes in approach/avoidance tendencies in individual patients. We assessed 24 PD patients on and off dopaminergic medication and 24 healthy controls (HC) performing a probabilistic reinforcement learning task, while undergoing functional magnetic resonance imaging. During learning, medication in PD reduced an overemphasis on negative outcomes. When patients were on medication, learning rates were lower for negative (but not positive) outcomes and concurrent striatal BOLD responses showed reduced prediction error sensitivity. Medication-induced shifts in negative learning rates were predictive of changes in approach/avoidance choice patterns after learning, and these changes were accompanied by striatal BOLD response alterations. These findings highlight dopamine-driven learning differences in PD and provide new insight into how changes in learning impact the transfer of learned value to approach/avoidance responses in novel contexts.

## INTRODUCTION

Learning from trial and error is a core adaptive mechanism in decision-making (Glimcher, 2002; Packard et al., 1989). This learning process is driven by reward prediction errors (RPEs) that signal the difference between expected and actual outcomes (Houk, 1995; Montague et al., 1996; Schultz et al., 1997). Substantia nigra (SN) and ventral tegmental area (VTA) midbrain neurons use bursts and dips in dopaminergic signaling to relay positive and negative RPE to prefrontal cortex (Deniau et al., 1980; Swanson, 1982) and the striatum, activating the so-called Go and NoGo pathways (Beckstead et al., 1979; Surmeier et al., 2007).

PD is caused by a substantial loss of dopaminergic neurons in the SN (Edwards et al., 2008), leading to the depletion of dopamine in the striatum (Koller and Melamed, 2007). Dopaminergic medication has been shown to alter how PD patients learn from feedback (Bódi et al., 2009; Cools et al., 2001) and how they use past learning to make choices in novel situations (Frank, 2007; Frank et al., 2004; Shiner et al., 2012). A common finding is that, after learning, when patients are on compared to off medication, they are better at choosing the option associated with the highest value (approach), whereas when off medication, they are better at avoiding the option with the lowest value (avoidance) (Frank, 2007; Frank et al., 2004). However, it is currently unknown how dopamine-induced changes during the learning process relate to these subsequent dopamine-induced changes in approach/avoid choice behavior on a within-patient basis.

An influential framework of dopamine function in the basal ganglia proposes that the dynamic range of phasic dopamine modulation in the striatum, in combination with tonic baseline dopamine levels, gives rise to the medication differences observed in PD (Frank, 2005). This theory suggests that lower baseline dopamine levels in unmedicated PD are favorable for the upregulation of the NoGo pathway, leading to an emphasis on learning from negative outcomes. In contrast, higher tonic dopamine levels in medicated PD lead to continued suppression of the NoGo pathway, resulting in (erroneous) response perseveration even after negative feedback. Extremes in these medication-induced changes in brain signaling are thought to manifest behaviorally in dopamine dysregulation syndrome, in which patients exhibit compulsive tendencies, such as pathological gambling or shopping (Voon et al., 2010). In support of the theory on Go/NoGo signaling, impairments in learning performance associated with higher dopamine levels have been found mainly in negative-outcome contexts; during probabilistic selection (Frank et al., 2004), reversal learning (Cools et al., 2006), and probabilistic classification (Bódi et al., 2009).

In addition to these behavioral adaptations, increased striatal activations have been reported in medicated PD patients during the processing of negative RPE (Voon et al., 2010). Similarly, a recent study on rats performing a reversal learning task revealed a distinct impairment in the processing of negative RPE with increased dopamine level (Verharen et al., 2018). However, very little is known about how these medication-related changes in the striatum’s responsivity to RPE relate to 1) later behavioral choice patterns, and 2) changes in brain activity during subsequent value-based choices.

We examined the role of dopaminergic transmission not only in choice behavior, but also in terms of associated neural mechanisms. 24 PD patients ON and OFF dopaminergic medication and a reference group of 24 age-matched HC performed a two-stage probabilistic selection task (Frank et al., 2004) (Figure 1A) while undergoing functional magnetic resonance imaging (fMRI). The experiment’s first stage was a learning phase, during which participants gradually learned to make better choices for three fixed pairs of stimulus options, based on reward feedback. In the second, transfer stage, participants used their learning phase experience to guide choices when presented with novel combinations of options, without receiving any further feedback (Figure 1A). Value-based decisions during the transfer phase were examined using an approach/avoidance framework (Figure 1B). To better describe the underlying processes that contribute to learning, behavioral responses were fit using a hierarchical Bayesian reinforcement learning model (Jahfari et al., 2017; van Slooten et al., 2018), adapted to estimate both within-patient effects of medication and across-subject effects of disease (Sharp et al., 2016). This quantification of behavior then informed our model-based fMRI analysis, in which we examined medication-related changes in blood oxygen level dependent (BOLD) brain signals in response to RPEs during learning, as well as medication-related changes in approach/avoidance behavior and brain responses during subsequent value-based choices.

**Figure 1.**
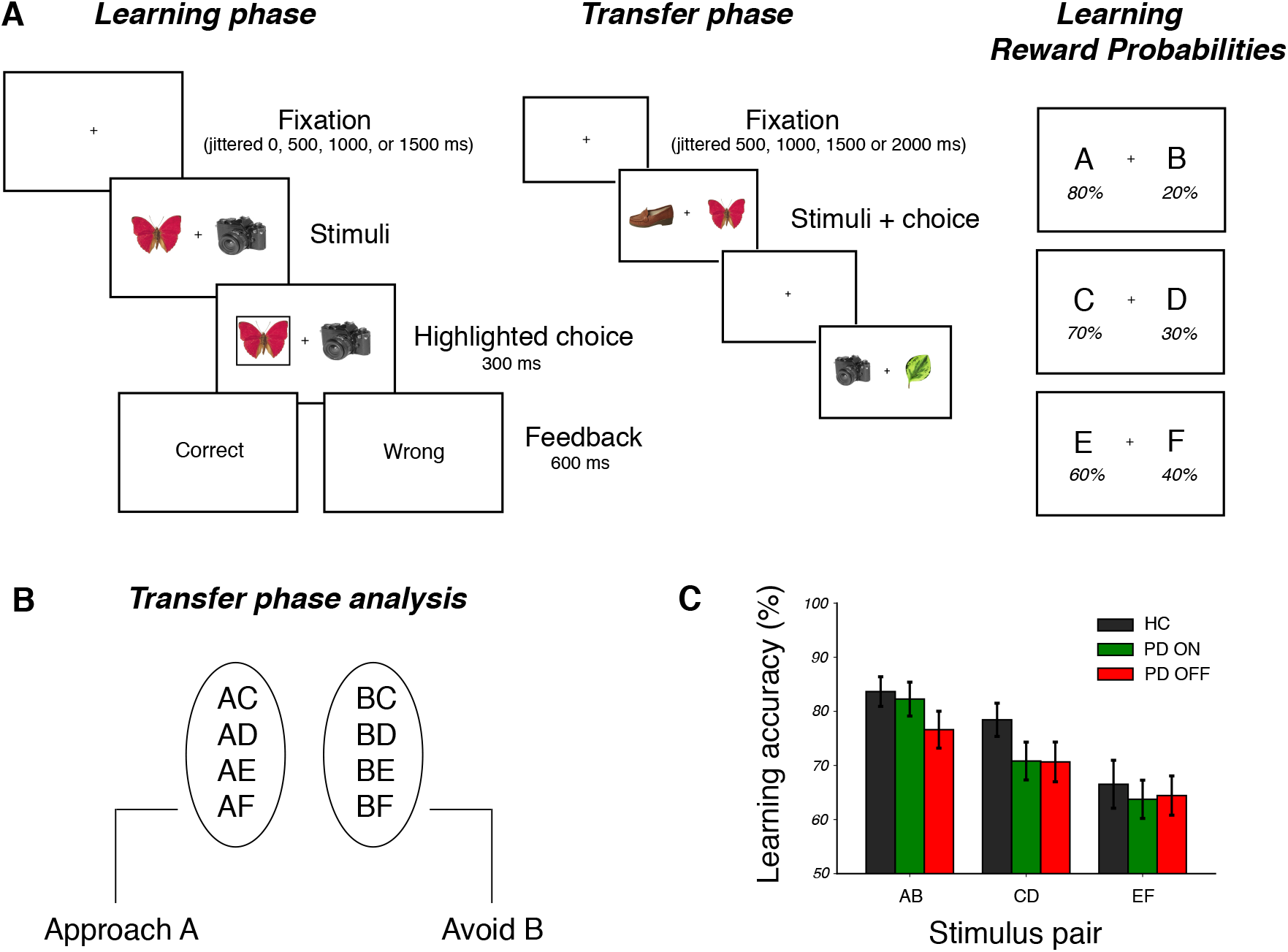
Learning and transfer phase tasks. **A)** *Learning task*: on each trial participants chose between two everyday objects and observed a probabilistic outcome ‘Correct’ or ‘Wrong’, corresponding to winning 10 cents or nothing. Each participant viewed three fixed pairs of stimuli (AB, CD, and EF) and tried to learn which was the best option of each pair, based on the feedback received. Reward probability contingency per stimulus during learning is shown on the right. *Transfer task*: participants were presented with all possible combinations of stimuli from the learning phase and had to choose what they thought was the better option, based on what they learned. No feedback was provided in this phase. **B)** Transfer phase analysis was performed on correctly choosing A on trials in which A was paired with another stimulus (approach accuracy) or correctly avoiding B on trials where B was paired with another stimulus (avoidance accuracy). **C)** Accuracy in choosing the better option of each pair across each group during learning (mean ±1 SEM). Parameter estimates of these medication and disease effects are presented in Figure S1.

## RESULTS

During the learning phase, participants successfully learned to choose the best option out of three fixed pairs of stimuli (Figure 1C). Each pair was associated with its own relative reward probability among the two options, labeled as AB (with 80:20 reward probability for A:B stimuli), CD (70:30) and EF (60:40). Choice accuracy analysis showed that learning took place in PD OFF, PD ON and HC (N=23 in each condition), with the probability with which participants chose the better option of each stimulus pair largely reflecting the underlying reward probabilities (PD ON: 82.3 % ± 3.1, 70.8 105 % ± 3.5, and 63.7 % ± 3.5; PD OFF: 76.6 % ± 3.4, 70.7 % ± 3.7, and 64.4 % ± 3.6; and HC: 83.7 % ± 2.7, 78.4 % ± 3.1, and 66.5 % ± 4.4 for AB, CD, and EF stimulus pairs, respectively).

We detailed within- and between-subject differences in choice accuracy using a Bayesian mixed-effects logistic regression on the observed trial-by-trial behaviors (see Methods). This analysis assessed how choice accuracy was affected by stimulus pair, medication, disease status, and their interactions. When patients were ON medication, overall performance was more accurate in comparison to OFF, with the biggest difference for the easier AB choices and relatively smaller difference for the more uncertain EF pair. This was evidenced by a main effect of stimulus pair (β (SE) = 0.35 (0.03), *z =* 10.19, *p* << .001), medication (β (SE) = 0.11 (0.04), *z* = 2.80, *p* = .005), and specifically, an interaction between medication and stimulus pair (β (SE) = 0.17 (0.05), *z* = 3.47, *p* <.001). Importantly, this specific effect of medication was reflected in a similar effect of disease when comparing PD OFF to HC, with a significant interaction between disease status and stimulus pair (β (SE) = 0.20 (0.05), *z* = 3.81, *p* < .001). Overall, these first analyses first analyses show an improvement in choice accuracy when patients are ON compared to OFF medication, with performance on the easiest option pair elevated up to the level of HC. However, although choice accuracy provides us with a general assessment of medication effects on performance, it does not relate these effects to a mechanistic explanation of how underlying indices of learning might be affected by medication. These underlying mechanisms can be studied and defined both at the group level (HC vs. PD), and within-subject level (PD ON vs. OFF) by adopting a formal learning model of behavior, to which we turn next.

### Medication reduces learning rate for negative outcomes

Reinforcement learning theories describe how an agent learns to select the highest-value action for a given decision, based on the incorporation of received rewards (Rescorla and Wagner, 1972; Sutton and Barto, 1998). We implemented a Q-learning model, graphically represented in Figure 2A, to describe both value-based decision-making and the integration of reward feedback in our experiment (Daw et al., 2011; Jocham et al., 2011; Schmidt et al., 2014). Our model used separate parameters to describe, for a given agent, how strongly current value estimates are updated by positive (α_gain_) and negative (α_loss_) feedback, i.e. positive and negative learning rates (Grogan et al., 2017; Jahfari et al., 2017; Slooten et al., 2018; Verharen et al., 2018), as well as a parameter that determines the extent to which differences in value between stimuli are exploited (β). To understand how medication affects learning in PD we examined the posterior distributions of group-level parameters representing the within-subject medication shift in α_gain_, α_loss_ and β (Figure 2B). The large leftward shift of the α_loss_ posterior distribution indicates higher learning rates after negative outcomes in PD OFF compared to ON (Bayes Factor (*BF*) = 11.40). This is consistent with the theory that PD increases the sensitivity to negative outcomes, and that dopaminergic medication remediates specifically this disease symptom. Conversely, shifts in the distributions of the α_gain_ and β parameters were merely anecdotal (1<*BF*s<2, see Table S4, and Figure S3 for individual results). For parameter comparisons based on disease status, we found strong evidence for a higher β i.e. greater exploitation, in HC compared to PD (BF = 16.89; see Figure S4).

**Figure 2.**
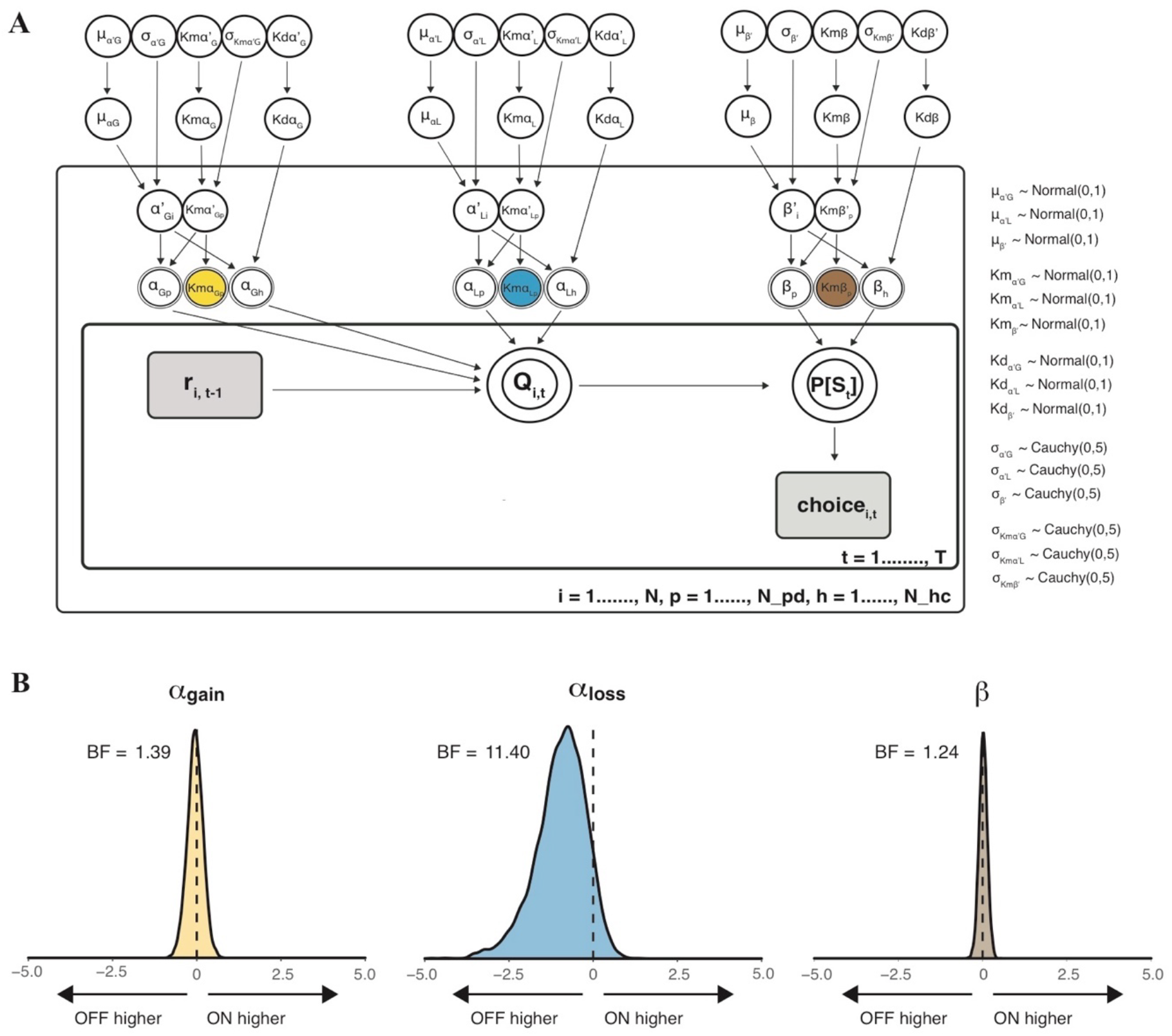
Q-learning graphical model and medication-driven parameter differences in PD. **(a)** Graphical representation of the hierarchical Bayesian Q-learning model. The inner box represents trial-by-trial information (*t*=1,…,T) and behavior of each subject. The outer box contains subject-level positive and negative learning rate (α_G_ and α_L_) and explore-exploit (β) parameters, for any *i*th subject (*i*= 1,…,N), *p*th PD subject (*p*=1,…,N_pd), or *h*th HC subject (*h*=1,…,N_hc), as well as medication (*Km)* and disease (*Kd)* difference variables for these parameters. The outermost layer represents group mean and standard deviation of each parameter across all subjects, medication difference distributions and disease difference distributions. Directed arrows from parents to children represent that parameters of the child are distributed according to its parents; the priors of these distributions are listed to the right. Double-lined borders denote deterministic variables. Continues variables are represented by circular nodes, and discrete variables by square nodes. Observed variables are shaded in grey. Per subject and session, *r_i,t-1_* is the reward received on the previous trial of a particular option pair, *Q_i,t_* is the current expected value of a particular stimulus, *P[S]* is the probability of choosing a particular stimulus in the current trial. **(b)** Group-level PD medication (within-subject) difference in learning phase parameters from Q-learning model (associated group-level *Km* medication difference parameters are highlighted in **a**). A leftward shift in the α_loss_ distribution indicates greater learning from negative outcomes in PD OFF compared to ON.

### Medication in PD reduces the sensitivity of dorsal striatum to RPE

In the Q-learning model, the learning rate weighs the extent to which value beliefs are updated based on trial-by-trial RPE. The processing of choice outcomes is known to influence BOLD signals in the striatum, where the sensitivity to RPE is changed when dopamine levels are manipulated (Jocham et al., 2011; Pessiglione et al., 2006; Schmidt et al., 2014). To establish whether RPE processing in the current study was influenced by dopaminergic state, we first examined within-subject medication-related differences in whole-brain responses to all positive and negative RPE in the learning phase using a single-trial GLM (see Methods). We found a significant PD OFF > ON medication difference in RPE modulation of the caudate nucleus and putamen (Figure 3), among other regions including the globus pallidus interna and externa, thalamus, cerebellum, lingual gyrus and precuneus. Because our model-based behavioral analysis revealed a medication-related difference specific to learning from negative outcomes (Figure 2B), we proceeded by analysing BOLD response time series to positive and negative outcomes separately.

**Figure 3.**
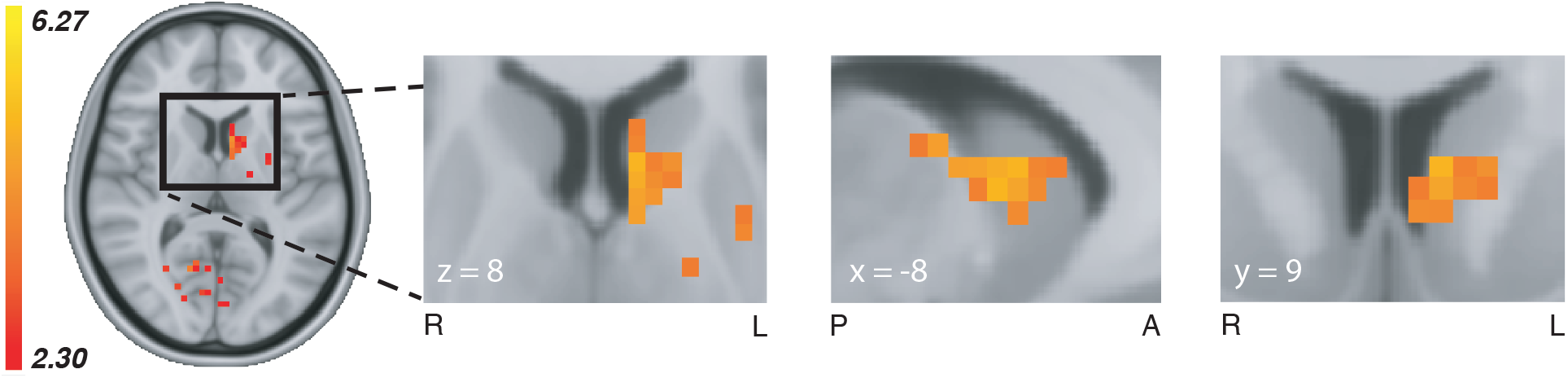
Whole-brain medication difference in RPE modulation. Whole-brain medication effects for the comparison PD OFF > ON in RPE-related modulations during the learning phase (*z*=2.3, *p*<.01, cluster-corrected), showing a dopamine-driven difference in the left dorsal striatum (see **Supplementary Table 5** for a full list of included brain areas and contrast statistics). Whole-brain group-level contrasts of RPE and feedback valence are available to view at https://doi.org/10.6084/m9.figshare.6989024.v1.

### Medication effects in dorsal striatum are specific to the processing of negative RPE

To disentangle the separate effects of positive and negative RPE signaling, we examined feedback-triggered BOLD time courses from three independent striatal masks; the caudate nucleus, putamen, and nucleus accumbens (see Methods). We found a significant medication difference only in the caudate nucleus, in BOLD activity associated only with negative RPE (Figure 4). RPE modulation was greater in PD OFF compared to ON, during the interval 7.51s – 10.67s after the onset of negative feedback. Medication status did not alter the BOLD responses to positive RPE, indicating that changes due to dopaminergic medication are specific to negative RPE signaling in the caudate nucleus, the most dorsal part of the striatum.

**Figure 4.**
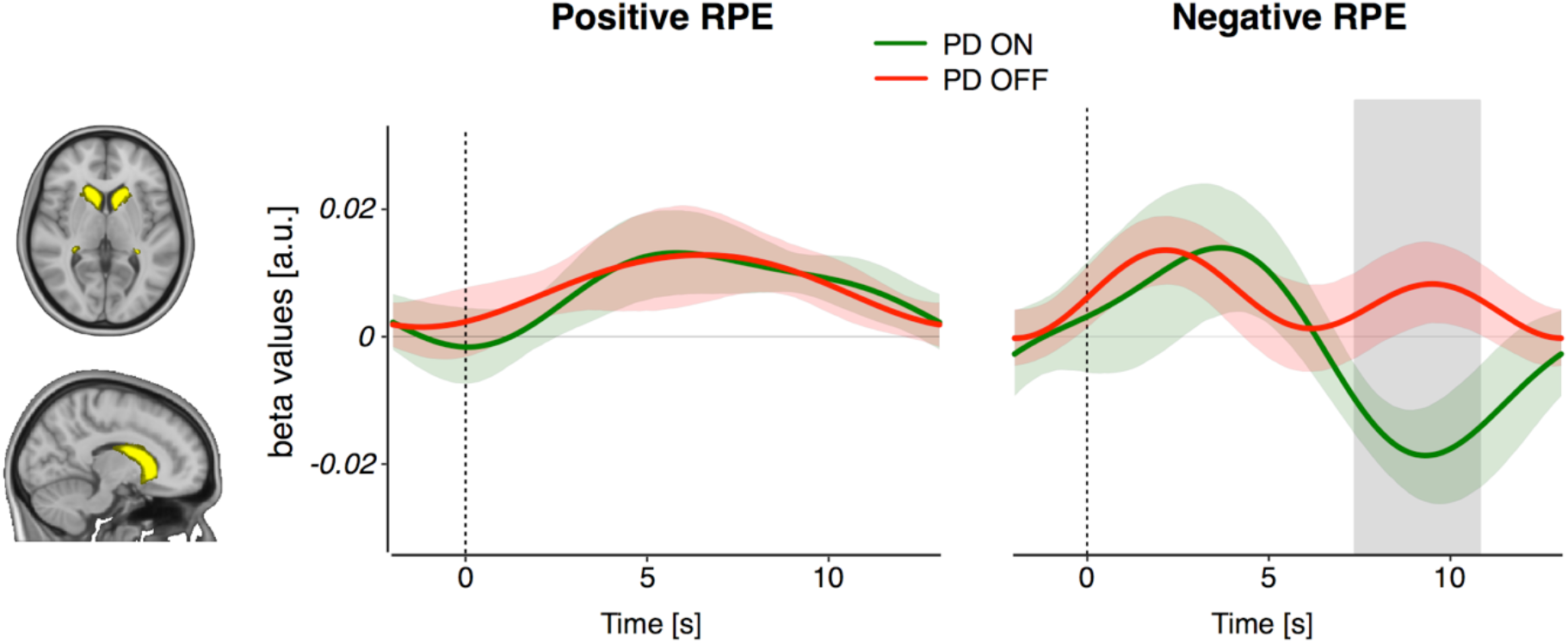
Positive and negative RPE modulations of the caudate nucleus. BOLD RPE covariation time courses for positive (left panel) and negative (right panel) feedback events. We found a significant difference between PD OFF and ON in negative RPE responses, but not in positive RPE responses. The grey shaded area reflects a significant PD OFF > ON difference passing cluster-correction for multiple comparisons across timepoints (p<.05). Colored bands represent 68% confidence intervals (±1 SEM). The same analysis of putamen and nucleus accumbens ROIs revealed no medication-related RPE differences (see Figure S6). Time courses of associated BOLD percent signal change after positive and negative outcomes in these striatal ROIs are presented in Figure S7.

### Behavioral analysis of transfer phase

The previous sections reveal how medication remediates the way patients learn from negative outcomes by detailing medication-related changes in brain and behavior. Much of the previous literature, however, has focused on how subsequent decision-making in the transfer phase is affected by dopaminergic medication (Frank, 2007; Frank et al., 2004; Grogan et al., 2017; Shiner et al., 2012). In the transfer phase, participants were presented with novel pairings of the learning phase stimuli and were asked to choose the best option based on their previous experience with the options (Figure 1A). We examined accuracy in correctly choosing the stimulus associated with the highest value from the learning phase (“Approach A”) and correctly avoiding the stimulus associated with the lowest value (“Avoid B”) (Frank, 2007; Frank et al., 2004; Grogan et al., 2017; Jocham et al., 2011; Shiner et al., 2012), as in Figure 1B (also see Methods). Consistent with several previous reports (Frank, 2007; Frank et al., 2004), results showed a strong interaction between medication (PD ON or OFF) and trial type (Approach A or Avoid B) (β (SE) = 0.34 (0.06), *z* = 5.75, *p* < .001). That is, medication in PD improved accuracy scores for “approach” trials, but decreased accuracy for “avoid" trials (Figure 5A, and see Figure S8 for HC performance). Notably, there were no main effects of trial type, medication or disease status in addition to this pivotal approach/avoid-medication interaction. Thus, medication only influenced Approach A versus Avoid B choice patterns, with no further differences in the overall accuracy across groups or trials.

### Medication shifts in learning rate for negative outcomes relate to behavioral and striatal changes during transfer

We have described how medication affects patients’ updating of their beliefs after encounters with negative feedback, and replicate previous work by showing medication-induced changes in approach/avoidance choices during a follow-up transfer phase with no feedback. This final section explores how the shift in learning rates caused by medication during learning relates to the subsequent approach/avoidance interaction in 1) choice outcomes, and 2) the BOLD response of the dorsal striatum.

Consistent with the observation that medication only affects learning rates after negative outcomes, we found that only the medication-related shift in α_loss_ (and not α_gain_) was predictive for the magnitude of change in approach/avoid behavior, as indicated by the lowest BIC in a formal model comparison analysis (see Methods and Table S6). In other words, the more α_loss_ was lowered by medication, the bigger the medication-induced interaction effect in future approach/avoidance choice patterns (β (SE) = 91.97 (41.26), *t* (22) = 2.23, *p* = .037, Figure 5B).

Because the dorsal striatum was differentially responsive to RPE during learning, we additionally explored how learning rate shifts relate to the striatal BOLD response in approach/avoidance trials, while patients were ON or OFF medication. To this end, we masked the caudate and putamen using the whole-brain RPE z-statistics map shown in Figure 3. From these masks BOLD responses were extracted for Approach A and Avoid B trials, for each of the PD ON and OFF sessions. Again, only the medication-induced shift in α_loss_ predicted the magnitude of change in the BOLD response of the caudate nucleus, but not the putamen, for approach/avoidance trials of OFF compared to ON (see Methods and Table S5) (β (SE) = 1.54, (0.56), *t* (22) = 2.77, *p*=.012, Figure 5C). In summary, these findings show that within-subject medication-related shifts in learning from negative outcomes are predictive of subsequent approach/avoidance medication-related changes, both in terms of behavioral accuracy and BOLD signaling in the caudate nucleus.

**Figure 5.**
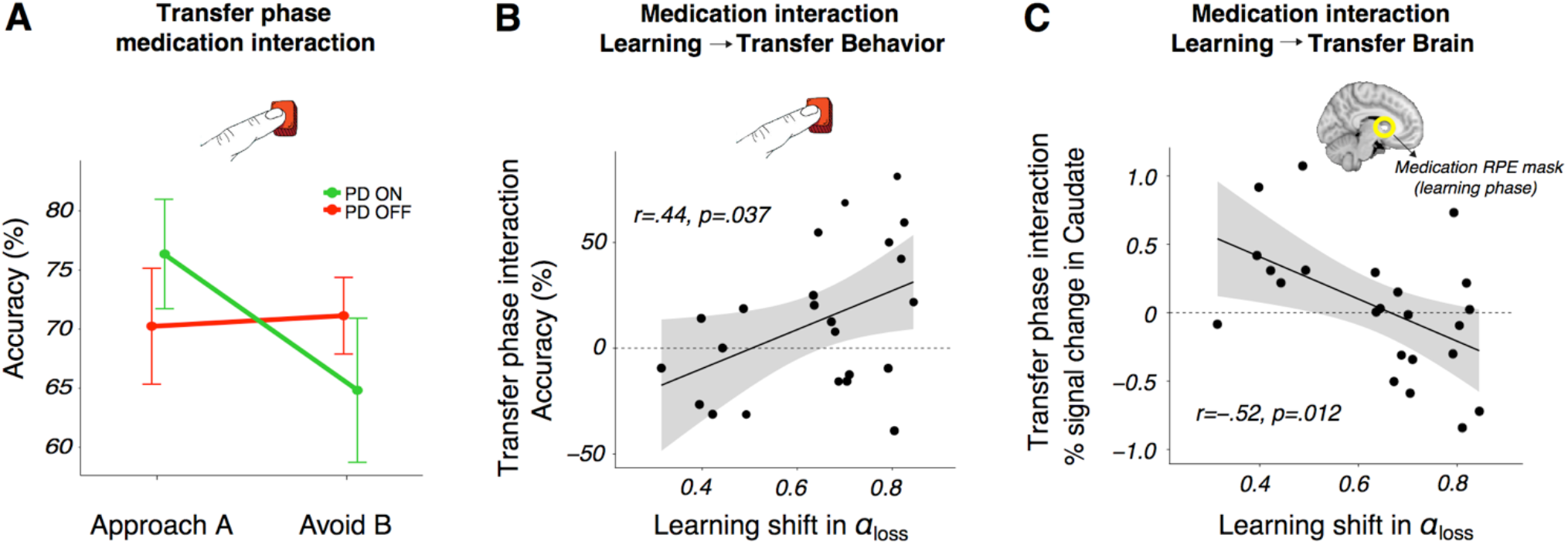
Medication-induced changes in learning from negative outcomes predicts the magnitude of medication difference in subsequent approach/avoidance behavioral choices and striatal response. **A)** Transfer phase behavioral accuracy in Approach A and Avoid B responses, showing a significant medication interaction in approach/avoidance behavior (*p*<.001). PD ON had a higher accuracy in approach trials but a lower accuracy in avoid trials than PD OFF. **B)** A positive relationship between the medication difference (i.e. the parameter shift for OFF > ON) in negative learning rate and the transfer phase medication accuracy difference (OFF > ON) in avoiding the lowest-value stimulus versus approaching the highest-valued stimulus, i.e. the interaction observed in **A**. **C)** A negative relationship between the medication difference (OFF > ON) in negative learning rate and the same transfer phase medication difference (OFF > ON) in avoid versus approach trials, here in terms of BOLD percent signal change in the caudate nucleus.

## DISCUSSION

Our findings provide a bridge between a previously disparate set of findings relating to reinforcement learning in PD. First, using a formalized learning theory, we show how dopaminergic medication remediates learning behavior by reducing the patient’s emphasis on negative outcomes. These behavioral adaptations were tied to BOLD changes in the dorsal striatum, with medication reducing the sensitivity to RPE, specifically during the processing of negative outcomes. Second, we explored how the medication-induced change in learning relates to subsequent approach/avoid choices that differ in PD when patients are ON or OFF medication. We found that the greater the degree of restoration by medication in the learning rate for negative outcomes, the greater the medication-related impact on both subsequent behavior and associated BOLD responses of the dorsal striatum during approach and avoidance choices.

Our finding that medication reduces negative learning rate directly replicates studies showing a medication-driven impairment in behavioral responses relating to negative feedback, in a variety of probabilistic learning tasks (Bódi et al., 2009; Cools et al., 2006; Frank et al., 2004; Palminteri et al., 2009). Furthermore, this finding corroborates a dopamine-driven reduction in model-based negative learning rate in PD patients (Voon et al., 2010) and rats (Verharen et al., 2018). The shift towards lower sensitivity to negative outcomes in PD ON reflects a partially restorative effect, as sensitivity to negative outcomes became more similar to that observed in healthy controls, whereas decision-making volatility, i.e. the exploitation of higher-valued options, did not (see Figure S5).

The medication interaction in subsequent approach/avoidance behavior we find in the transfer phase supports previous research on the transfer of learned value to new contexts (Cox et al., 2015; Frank, 2007; Frank et al., 2004). This reinforces the notion that dopaminergic medication shifts the balance in activation of the Go and NoGo pathways of the striatum (Frank, 2005). It has been an open question whether these Go and NoGo pathways are in competition with each other or function independently. A recent review suggests that the Go and NoGo pathways should not be viewed as separate, parallel systems (Calabresi et al., 2014). The two pathways are instead described to be structurally and functionally intertwined, with “cross-talk” occurring between Go and NoGo neuronal subtypes. This represents a potential means by which the dopamine-dependent alterations in learning from negative outcomes observed in the current study can lead to an integrated (interactive) effect on subsequent approach and avoidance behavior and associated BOLD activation in the striatum.

We observed greater RPE modulation of BOLD signaling in PD OFF compared to ON, indicating a medication-related role in the modulation of caudate nucleus activity during learning. Striatal BOLD activations have previously been demonstrated to track RPE, with numerous studies implicating the caudate nucleus in RPE signaling during goal-directed behavior (Davidson et al., 2004; Delgado et al., 2005; Haruno and Kawato, 2006; O’Doherty et al., 2004). The whole-brain analysis used in the current study reveals greater within-subject RPE modulation in patients OFF compared to ON medication in the dorsal striatum, a region well established to suffer substantial depletion of dopamine availability in PD (Bernheimer et al., 1973; Dauer and Przedborski, 2003). Patients in our study do not exhibit clear medication-related differences that signify an excessive level of dopamine in the ventral striatum, as postulated by the dopamine overdose hypothesis (Cools et al., 2006, 2001) and presented in studies focusing on the nucleus accumbens (Cools et al., 2007; Schmidt et al., 2014). In our data, there does appear to be a quantitative medication-induced increase in the modulation of nucleus accumbens activity by positive RPE, however, this effect is not significant (see Figure S6).

Activation of the dorsal striatum has been reported for instrumental but not Pavlovian learning, suggesting its role in establishing stimulus-response-outcome associations (O’Doherty et al., 2004). A prominent theory of dopamine functioning, the actor-critic model, highlights distinct roles for reward prediction and action-planning in reinforcement learning (Houk, 1995; Joel et al., 2002; Suri and Schultz, 1999), with the ventral striatum (critic) implicated in the prediction of future rewards (Cardinal et al., 2002), and the dorsal striatum (actor) proposed to maintain information about rewarding outcomes of current actions to help inform future actions (Atallah et al., 2007; Packard and Knowlton, 2002). Connectivity between the midbrain SN and dorsal striatum has also been found to predict the impact of differing reinforcements on future behavior (Kahnt et al., 2009). Overall, the caudate nucleus has been put forward as a hub that integrates information from reward and cognitive cortical areas in the development of strategic action planning (Haber and Knutson, 2010). The dopamine-dependent differences in RPE modulation of BOLD activity in dorsal striatum presented here therefore suggest that PD’s dopamine-related effects are specific to the processing of feedback to guide future actions. The dopamine-related interaction in approach/avoidance behavior found in the transfer phase, in which actions were guided by previously learned values, provides further support for this interpretation.

We were able to link medication-dependent changes in learning from negative outcomes to subsequent changes in approach/avoidance striatal activity by specifically focusing on the region that showed a robust medication-dependent difference in phasic RPE modulation during learning. This suggests that the caudate nucleus’ processing of negative RPE in PD ON plays an important role in the subsequent medication-induced shift in balance between approach and avoidance behavior. Although focusing on the ventral striatum, a recent study on rats showed that increased activation in the VTA-NAc pathway associated with a higher dopaminergic state was reflected in behavior by a reduced sensitivity to negative outcomes (Verharen et al., 2018). Our findings suggest that the caudate nucleus may play a similar role in the processing of negative outcomes in PD. Future research could address whether this is modulated by SN-caudate nucleus connectivity and/or the interplay between instrumental and Pavlovian learning.

In several previous studies, dopamine level was manipulated pharmacologically in healthy adults, via levodopa medication (Pessiglione et al., 2006) or NoGo (D2) receptor antagonists (Jocham et al., 2011; Van Der Schaaf et al., 2014). Here, we examined separable disease-related and dopaminergic medication-related effects in PD. Patients in the current study used a combination of dopaminergic medications, including those acting on both Go and NoGo receptors (levodopa), inhibitors that slow the effect of levodopa to give a more stable release, and dopamine agonists, which have a particular affinity for NoGo receptors. Accordingly, a limitation of our study is that we cannot pin down the relationship between specific dopaminergic medications and changes in learning. Dissociation between the different types of dopaminergic medication could therefore be a potential avenue for future research.

Although there is moderate evidence for a higher sensitivity to negative feedback in PD OFF compared to HC (see Figure S5), we find that the greatest disease-related difference lies in the explore/exploit parameter of the model (Figure S4). Higher choice accuracy during easier decisions in HC is likely strongly influenced by greater exploitation of value differences between options; indeed, a positive correlation has recently been shown between choice accuracy and exploitation in a similar reinforcement learning task (Jahfari et al., 2017). In the current study, this difference in exploitation was observed regardless of PD medication state (Figure S5), showing that dopamine medication in PD does not reinstate healthy exploitative behavior. This selectivity of dopaminergic medication’s effects on learning may indicate certain mechanisms underlying PD-related psychiatric disorders (Voon et al., 2010). Recent evidence from a perceptual decision-making study in PD showed an impaired use of prior information in patients in making perceptual decisions (Perugini et al., 2016), a deficiency that also was not alleviated by dopaminergic medication (Perugini et al., 2018). Thus, regardless of medication status, PD patients show impairment in the integration of memory with the current sensory input. Since the explore/exploit parameter of the task used in our experiments is dependent upon the retrieval of the expected value of chosen options, a similar memory-guided decision-making impairment may have also played a role in the current reinforcement learning task.

In conclusion, we comprehensively illustrate how dopaminergic medication used in PD can help remediate sensitivity to negative outcomes, indicated by both changes in negative learning rate and the dorsal striatum’s response to negative RPE. Furthermore, we show how, when using experience garnered during learning to guide subsequent value-based decisions, these effects shift the balance of approach/avoidance behavior and associated striatal activation. Aside from explicating dopamine’s role in reinforcement learning and value-based decision-making, our findings open new avenues of treatment in PD and its associated psychiatric symptoms.

## METHODS

### Participants

Twenty-four patients with Parkinson’s disease (7 female, mean age = 63 ± 8.2 years old) were recruited via the VU medical center, Zaans medical center, and OLVG hospital in Amsterdam. All patients were diagnosed by a neurologist as having idiopathic PD according to the UK Parkinson’s Disease Society Brain Bank criteria. Twenty-four age-matched healthy controls (9 female, mean age = 60.3 ± 8.5 years old) were also recruited, via the PD patients (e.g. spouses, relatives) or from the local community. This study was approved by the Medical Ethical Review committee (METc) of the VU Medical Center, Amsterdam.

Prior to participation, all subjects were screened according to the following inclusion criteria: age range 40-75 years old, normal/corrected-to-normal vision, and a prior diagnosis of PD in patients. Exclusion criteria were no current psychotropic medication usage (other than DA/Parkinson-related medication in patients), no major somatic disorder or psychosis, no dementia diagnosis, and no history of head injury, stroke or any other neurological diseases. Patients were additionally not included if they took selective serotonin reuptake inhibitors (SSRIs) in order to primarily examine the effects of dopamine, since serotonin has also been implicated in learning mechanisms (Daw et al., 2002; den Ouden et al., 2013). At each session of the study, the severity of clinical symptoms was assessed according to the Hoehn and Yahr rating scale(Hoehn and Yahr, 1967) and the motor part of the Unified PD Rating Scale (UPDRS III; Fahn et al., 1987). Demographic and clinical data of the included participants can be seen in Table S1. Information on Parkinson-related medication per patient is available in Table S2.

All participants provided written informed consent and were paid a minimum of €100 (PD patients, three sessions) or €70 (HCs, two sessions) for participation. A reward bonus was additionally paid out based on performance during the reinforcement learning task (PD ON mean= €8.86 ± 0.99, PD OFF mean= €8.85 ± 1.00, HC mean= €9.34 ± 1.42, per learning run).

We excluded one PD patient (excessive falling asleep in scanner) and one HC (could not learn the task) from both learning and transfer phase behavior and fMRI analyses. fMRI data of one further HC could not be analyzed (T1 scan was not collected; session was terminated early due to claustrophobia). Transfer phase fMRI and behavior data were not collected for an additional HC due to early termination of scanning session (technical malfunction). Overall, we included 23 PD patients on and off dopaminergic medication in all behavioral and fMRI analyses. 23 HCs were included in learning phase behavioral analysis, 22 in learning phase fMRI analysis, and 21 in transfer phase behavioral and fMRI analysis.

### Procedure

The study was set up as a dopaminergic-manipulation, within-subject design in PD patients, to reduce the variance associated with inter-individual differences. All PD and HC participants took part in at least two sessions, the first of which was always a neuropsychological examination (2 hours; 30 minutes of which was spent practicing the reinforcement learning task with basic shape stimuli). PD patients subsequently attended two separate fMRI scanning sessions (once in a dopamine-medicated “ON” state and once in a lower dopamine “OFF” state), and HCs underwent one fMRI session. The patient fMRI sessions were carried out over the same weekend in all but one patient (2 weeks apart) and were counterbalanced for medication/order effect in the following way: first session ON and second session OFF in 12 patients, with the opposite schedule in the remaining 12 patients. All OFF sessions had to be carried out in the morning for ethical reasons. Patients were instructed to not take their usual dopamine medication dosage on the evening prior to and the morning of the OFF session, thereby allowing >12 hours withdrawal at the time of scanning. If patients were on dopamine-agonists (pramipexole, ropinerol) they took their final dopamine-agonist dose on the morning prior to the day of scanning (~ 24 hours withdrawal). One PD patient took his medication 8.5 hours before OFF day scanning to relieve symptoms but was still included in the analysis.

### Neuropsychological assessment

Participants completed a battery of neuropsychological tests on their first visit. A description of these tests and self-report questionnaires, along with group results, is included in Table S1. All patients used their dopaminergic medication as usual during this session. These assessments were not examined in the current study, but are discussed in greater detail elsewhere (Engels *et al.*, 2018).

### Reinforcement Learning Task

Participants completed a probabilistic selection reinforcement learning task consisting of two stages; a learning phase and transfer phase. This task has been used in a number of previous studies, in both PD patients (Frank et al., 2004; Grogan et al., 2017; Shiner et al., 2012) and healthy participants (Jocham et al. 2011; Jahfari et al. 2017; van Slooten et al. 2018). We used pictures of everyday objects from different object categories, such as hats, cameras, and leaves (stimulus set extracted from Konkle *et al.*, 2010, 17-objects.zip).

### Learning phase

In the learning phase, three different pairs of object stimuli (denoted as AB, CD and EF) were repeatedly presented in random order. Each pair had assigned reward probabilities associated with each stimulus, and participants had to learn to choose the best option of each pair based on the feedback provided (see Figure 1A). Participants were instructed to try to find the better option of a pair in order to maximize reward. Feedback was either “Goed” or “Fout” text (meaning “correct” or “wrong” in Dutch), indicating a payout of 10 cents for correct trials and nothing for incorrect trials. Different objects were used across each fMRI session of patients, so as not to induce any familiarity or reward associations with particular stimuli. In the “easiest” AB pair, the probability of receiving reward was 80% for the A stimulus and 20% for the B stimulus, with ratios of 70:30 for CD and 60:40 for EF. The EF pair was therefore the hardest to learn due to a much closer reward probability between the two options. All object stimuli were counterbalanced for reward probability pair across subjects (i.e. a leaf and hat as the A and B stimuli for one participant were the C and D stimuli for another participant) and for better versus worse option of a pair (i.e. a leaf and hat as the A and B stimuli for one participant were the B and A stimuli for another participant). In total, there were 12 object stimuli and a single participant viewed six of these objects in a given fMRI session, with PD patients viewing the remaining 6 stimuli in their second fMRI session. The learning phase consisted of two runs of 150 trials each (50 trials per stimulus pair). Each run was interspersed with 15 null trials to minimize problems relating to temporal autocorrelation and model fitting of rapid event-related designs. Null trials, during which only the fixation cross was presented, lasted at least 4 seconds plus an additional interval generated randomly from an exponential distribution with a mean of 2 seconds. Each task trial had a fixed duration of 5000 ms, and began with a jittered interval of 0, 500, 1000, or 1500 ms to obtain an interpolated temporal resolution of 500 ms. During the interval, a black fixation cross was presented and participants were asked to hold fixation. Two objects were then presented simultaneously left and right (counterbalanced across left/right locations per pair) of the fixation cross and remained on the screen until a response was made. If a response was given on time, a black frame surrounding the chosen object was shown (300 ms) and followed by feedback (600 ms). Omissions were followed by the text “te langzaam” (“too slow” in Dutch). The fixation cross was displayed alone after feedback was presented, until the full trial duration was reached.

### Transfer phase

In the transfer phase, novel pairings of all possible combinations of the six stimuli were presented, in addition to the original three stimulus pairs, making up 15 possible pairings. This phase consisted of two runs of 120 trials each (8 trials per pair), and each run randomly interspersed with 12 null trials. The duration of these null trials was generated in the same way as in the learning phase. Participants were instructed to choose what they thought was the better option, given what they had learned. There was no feedback in this phase and no frame surrounding the chosen response. Each trial began with a jittered interval of 500, 1000, 1500 or 2000 ms, with a new trial starting whenever a response was made.

Each object stimulus was presented equally often on the left or right side in both learning and transfer phases. Responses were made with the right hand, using the index or middle finger to choose the left or right stimulus, respectively. One patient felt uncomfortable using two fingers of the right hand and so responded with his left and right index finger on separate button boxes (in both ON and OFF sessions). The feedback text was made larger for one patient in both ON and OFF sessions to make it easier to read.

### Behavioral analysis

#### Learning phase

Bayesian mixed-effects logistic regression modeling was carried out on trial-by-trial behavior (Doll et al., 2016; Sharp et al., 2016; Wunderlich et al., 2012). These analyses were performed in R (R Development Core and Team, 2017), using the Bayesian Linear Mixed-Effects Models (blme) package (Chung et al., 2013), built on top of lme4 (Bates et al., 2014). In our mixed-effects models we coded for both fixed and random trial-by-trial effects and allowed for a varying intercept on a per subject basis. For the model on learning behavior, the dependent variable was accuracy in choosing the better stimulus of a pair (correct=1, incorrect=0). Stimulus pair was taken as a within-subject (random-effect) explanatory variable (EV), from easiest to most difficult (AB pair=1, CD pair=0, EF pair=-1). We also included two binary covariates, which interacted with the within-subject random effects variable: the between-subject effect of disease (PD=0, HC=1) and the within-subject effect of dopaminergic medication state (OFF=0, ON=1). HCs were also coded as OFF=0, as we wanted this variable to capture only the within-subject effect of medication. Since disease and medication status were both included in the same model, PD OFF was considered to act as a baseline (Dis=0, Med=0). Within-subject effects of medication for PD ON (Dis=0, Med=1) were therefore captured by the medication variable only and between-subject effects of disease for HC (Dis=1, Med=0) were captured by the disease variable only (with Dis=1 essentially meaning “healthy”). Positive beta estimates obtained from the model therefore indicated higher accuracy for either PD ON or HC compared to OFF in the Med and Dis variables, respectively, with negative estimates for those variables reflecting greater accuracy for PD OFF.

#### Transfer phase

The mixed-effects regression on transfer phase behavior was carried out on trials in which either the A or B stimulus appeared, excluding those in which both appeared together (see Figure 1B). The expectation was that participants should opt to choose A (Approach A) and avoid choosing B (Avoid B) whenever they were presented, since they were associated with the highest and lowest reward probabilities during learning, respectively. The regression was performed almost identically to that in the learning phase, except the stimulus pair variable was replaced with an Approach A / Avoid B trial variable (A=1, B=-1). The dependent variable (accuracy) was then coded as 1 for correctly choosing A in Approach A or correctly not choosing B in Avoid B trials, and as 0 for incorrectly choosing the other option each trial type.

To further examine the medication difference found in Approach A vs. Avoid B accuracy of the transfer phase (Figure 5A), we performed robust (multiple) regression with OFF > ON medication difference in Avoid B > Approach A accuracy as the dependent variable and compared three models with within-subject learning rate medication differences derived from the learning phase Q-learning model as EVs. The EVs were either: both positive and negative learning rate medication differences (termed kα_gain_ and kα_loss_), kα_gain_ only, or kα_loss_ only. BIC values were calculated and compared across these models (see Table S6). The learning to transfer phase accuracy correlation *p*-value was therefore obtained from the winning α_loss_-only model. Individual medication differences were quantified as the modes of the within-subject medication difference parameter distributions, to capture peak probability densities (see Figure 2A and *Group-level model*).

#### Computational model

We fit a hierarchical Bayesian temporal difference Q-learning model to learning phase behavioral data. The Q-learning reinforcement learning algorithm (Sutton and Barto, 1998) captures trial-by-trial updates in the expected value of options and has been used extensively to model behavior during learning (Daw et al., 2011; Grogan et al., 2017; Jahfari et al., 2017; Jocham et al., 2011; Schmidt et al., 2014). We used a variant of this model with three parameters free to vary across subjects, allowing us to determine how subjects learned separately from positive and negative feedback (α_gain_ and α_loss_) and how much they exploited differences in value between stimulus pair options (β). In hierarchical models, group and individual parameter distributions are fit simultaneously and can constrain each other, leading to greater statistical power over standard non-hierarchical methods (Ahn et al., 2011; Jahfari et al., 2017; Kruschke, 2015; Steingroever et al., 2013; Wiecki et al., 2013). We also fit a separate model with an additional free parameter, relating to persistence of choices irrespective of feedback, and performed model comparison, to justify that the chosen model better represented the data (see Table S3).

#### Subject-level model

The Q-learning algorithm assumes that after receiving feedback on a given trial, subjects update their expected value of the chosen stimulus (0 < *Q_chosen_* < 1) based on the difference between reward received for choosing that stimulus (*r_chosen_,* reward or no reward) and their prior expected value of that stimulus, according to the following equation:

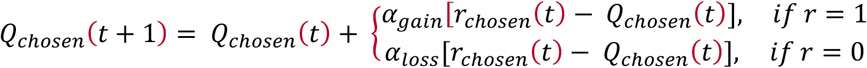

The term *r_chosen_*(*t*) − *Q_chosen_*(*t*) is the reward prediction error (RPE). Accordingly, choices followed by positive feedback (*r* = 1) are weighted by the α_gain_ learning rate parameter and choices followed by negative feedback (*r* = 0) are weighted by the α_loss_ learning rate parameter (0 < α_gain,_ α_loss_ <1). All Q-values were initialized at 0.5 (no initial bias in value). The probability of choosing one stimulus over another is described by the softmax rule:

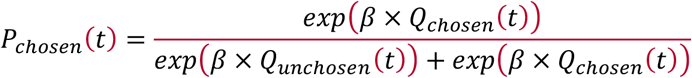

where β is known as the inverse temperature or “explore-exploit” parameter (0 < β < 100). Effectively, β is used as a weighting on the difference in value between the two options. The free parameters α_gain_, α_loss_ and β were fit for each subject individually, in a combination that maximizes the probability of the actual choices made by the subject.

#### Group-level model

The subject-level model described above was nested inside a group-level model in a hierarchical manner. All participants were assigned to one group and we fit separate group-level distributions to capture the within-subject effect of medication and the across-subject effect of disease (see model representation in Figure 2A), adopting a similar group-level model to that described in (Sharp et al., 2016). In Figure 2A, the free parameters α_gain_ and α_loss_ are denoted as α_G_ and α_L_ for viewing purposes. The inner box describes trial-by-trial behavior of individual subjects. The quantities r_i, t-1_ (reward for participant i on trial t-1) and choice_i, t-1_ (choice made by participant i on trial t-1) were obtained directly from the data. The quantities with a double-border were deterministic; e.g. Q-values (Q_i, t-1_) and the probability of selecting a certain stimulus (P[S_t_]) were calculated according to the equations above, given the fitted free parameters for a particular subject. Higher layers of the model represent within-subject parameters (separately for Kα_gain_, Kα_loss_ and Kβ) that capture the medication effects per subject, and group-level parameters capturing the group-level (within-subject) effect of medication and group-level (across-subject) effect of disease. These parameters were transformed during estimation using an approximation of the probit transformation, indicated by *z*’_i_. This is the inverse cumulative distribution function of the normal distribution. *Φ_approx_* represents the standard cumulative normal distribution.

Similar to Sharp et al., 2016, we tested for medication- or disease-related variation in the three parameters α_gain_, α_loss_ and β by including additional intermediate terms. For α_gain_ these were: *KmαG’p* (for the effect of medication *Km* on α_gain_ in PD patient *p*) and *KdαG’h* (for the effect of no disease *Kd* on α_gain_ in HC participant *h*), with the same terms for α_loss_ *(KmαL’p* and *KdαL’h)* and β (*Km*β*’p* and *Kd*β*’h)*. Symmetric boundaries for all *Km* and *Kd* distributions were used to constrain the model and assist with convergence (−5 < *Km, Kd* < 5). As in the logistic regression we took PD OFF as “baseline” by using two binary indicators: (′*on*′) = 0, and (′*healthy*′) = 0. PD ON was coded as (′*on*′) = 1, (′*healthy*′) = 0, and HC was coded as *I*(′*on*′) = 0, *I*(′*healthy*′) = 1. For subject *s* and medication condition *m*, the αG parameter of an individual subject was distributed as follows:

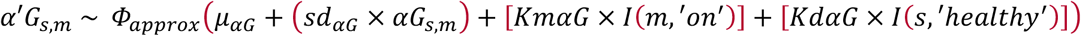

Using the binary indicators described above, PD OFF did not contain either of the or *KdαG* terms, PD ON included the *KdαG* term to indicate the within-subject effect of medication, and HC included the *KdαG* term to denote the between-subject effect of disease. α_loss_ and β parameters were distributed in the same way with their corresponding terms. Since the medication effect was within-subject, it was itself a subject-specific random variable with its own population-level mean and variance, given by:

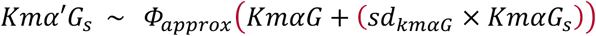

A normal prior was assigned to group-level means of the three free parameters and the *Km* and *Kd* indicators of each free parameters, *μ*_z’_ *N* (0,1). A half-Cauchy prior was given to all group-level standard deviations. σ_z’_ ~ *Cauchy* (0,5). Weakly informative priors such as these are recommended in small sample sizes to reduce the influence of the priors on posterior distributions (Ahn et al., 2017). 565 Bayes Factors (BF) of group level posterior distributions for medication and disease differences were calculated as the ratio of the posterior density above zero relative to the posterior density below zero (Pedersen et al., 2016). This method is possible since the priors for the distributions of these parameters were symmetric (unbiased) around zero (Marsman and Wagenmakers, 2017). Categories of evidential strength of an effect are based on (Jeffreys, 1998), with BFs > 10 considered as strong evidence that the shift in the posterior distribution is different from zero.

#### Model estimation

The model was estimated using Markov Chain Monte Carlo (MCMC) inference. Models were implemented using the Stan programming language (Hoffman and Gelman, 2016; Stan Development Team, 2014). We ran three chains of 5000 samples each (discarding the first 2500 of each chain for burn-in), and ensured convergence using manual examination of the trace plots (hairy caterpillars, easily moving around the parameter space) and evaluation of *r̂* statistics, which were all <1.1 (Gelman and Rubin, 1992). Simulations displayed in Figure S2 show adequate parameter recovery. Mean group-level posterior distributions after fitting can also be seen in Figure S2. We additionally generated a number of quantities of interest from the model, including individual subjects’ trial itself a subjects’ trial-by-trial RPE for inclusion in fMRI whole-brain and deconvolution analyses.

### fMRI image acquisition

fMRI scanning was carried out using a 3T GE Signa HDxT MRI scanner (General Electric, Milwaukee, WI, USA) with 8-channel head coil at the VU University Medical Center (Amsterdam, The Netherlands). Functional data for the learning and transfer phase runs were acquired using T2*-weighted echo-planar images with blood oxygenation level dependent (BOLD) contrasts, containing approximately 410 and 240 volumes for learning and transfer runs respectively. The first two TR volumes were removed to allow for scanner stabilization. Each volume contained 42 axial slices, with 3.3 mm in-plane resolution, TR = 2,150 ms, TE = 35 ms, FA = 80 degrees, FOV = 240 mm, 64 × 64 matrix. Structural images were acquired with a 3D T1-weighted magnetization prepared rapid gradient echo (MPRAGE) sequence with the following acquisition parameters: 1 mm isotropic resolution, 176 slices, repetition time (TR) = 8.2 ms, echo time (TE) = 3.2 ms, flip angle (FA) = 12 degrees, inversion time (TI) = 450 ms, 256 × 256 matrix. The subject’s head was stabilized using foam pads to reduce motion artifacts.

### fMRI analysis

#### Preprocessing

Preprocessing was performed using FMRIPREP version 1.0.0-rc2 (O. Esteban et al., 2018; Oscar Esteban et al., 2018), a Nipype-based tool (Gorgolewski et al., 2017, 2011). The following information was generated from FMRIPREP based on the preprocessing pipeline used in this study. Each T1w (T1-weighted) volume was corrected for INU (intensity non-uniformity) using N4BiasFieldCorrection v2.1.0 (Tustison et al., 2010) and skull-stripped using antsBrainExtraction.sh 601 v2.1.0 (using the OASIS template). Brain surfaces were reconstructed using recon-all from FreeSurfer 602 v6.0.1 (Dale et al., 1999), and the brain mask estimated previously was refined with a custom variation of the method to reconcile ANTs-derived and FreeSurfer-derived segmentations of the cortical gray-matter of Mindboggle (Klein et al., 2017). Spatial normalization to the ICBM 152 Nonlinear Asymmetrical template version 2009c (Fonov et al., 2009) was performed through nonlinear registration with the antsRegistration tool of ANTs v2.1.0 (Avants et al., 2008), using brain-extracted versions of both T1w volume and template. Brain tissue segmentation of cerebrospinal fluid (CSF), white-matter (WM) and gray-matter (GM) was performed on the brain-extracted T1w using fast (Zhang et al., 2001) (FSL version 5.0.9). Functional data was motion corrected using *mcflirt* (FSL version 5.0.9) (Jenkinson et al., 2002). "Fieldmap-less" distortion correction was performed by co-registering the functional image to the same-subject T1w image with intensity inverted (Huntenburg et al., 2012; Wang et al., 2017) constrained with an average fieldmap template (Treiber et al., 2016), 613 implemented with *antsRegistration* (ANTs). This was followed by co-registration to the corresponding T1w using boundary-based registration (Greve and Fischl, 2009) with 9 degrees of freedom, using *bbregister* (FreeSurfer version 6.0.1). Slice timing correction was not performed on the data. Motion correcting transformations, field distortion correcting warp, BOLD-to-T1w transformation and T1w-to-template (MNI) warp were concatenated and applied in a single step using *antsApplyTransforms* (ANTs version 2.1.0) using Lanczos interpolation. Physiological noise regressors were extracted applying CompCor (Behzadi et al., 2007). Principal components were estimated for the two CompCor variants: temporal (tCompCor) and anatomical (aCompCor). A mask to exclude signal with cortical origin was obtained by eroding the brain mask, ensuring it only contained subcortical structures. Six tCompCor components were then calculated including only the top 5% variable voxels within that subcortical mask. For aCompCor, six components were calculated within the intersection of the subcortical mask and the union of CSF and WM masks calculated in T1w space, after their projection to the native space of each functional run. Frame-wise displacement (Power et al., 2014) was calculated for each functional run using the implementation of Nipype. Many internal operations of FMRIPREP use Nilearn (Abraham et al., 2014), principally within the BOLD-processing workflow. For more pipeline details, please refer to https://fmriprep.readthedocs.io/en/latest/workflows.html.

#### Single-trial whole-brain analysis

We were interested in estimating trial-by-trial representations of learning (e.g. RPEs) in the brain. For this, we carried out a single-trial whole-brain analysis to capitalize on the variability in BOLD signal across trials, using Nipype’s FSL interface (Gorgolewski et al., 2017, 2011). A Least Squares All (LSA) GLM fit was performed on each subject’s brain data, per learning run (see Mumford et al., 2012). The feedback onset of each trial was included as a separate regressor. We included 13 confound regressors to remove nuisance effects that might have contributed to the brain signal: Framewise Displacement (FD), 6 rigid-body transform motion parameters (3 translational, 3 rotational), and 6 aCompCor physiological noise regressors (to help exclude physiological noise in the CSF and WM). Spatial smoothing was performed using a Gaussian kernel with a full width at half maximum of 4 mm. A 3^rd^ order Savitsky-Golay filter was used for high-pass filtering, with a window length of 120 seconds. The first-level design was set up using the Nipype interface to FMRI Expert Analysis Tool (FEAT) from FSL (FMRIB’s Software Library, www.fmrib.ox.ac.uk/fsl). Delta functions of all regressors in the model were convolved with the canonical hemodynamic response function (HRF) and regressed against each subject’s fMRI data, using Nipype’s FSL FILMGLS interface. From this, a contrast of parameter estimates (COPE) was obtained for each trial of every subject. Next, we performed a second-stage analysis on the single-trial copes, to model the per trial feedback valence and RPE. The feedback regressor was coded as +1 or −1 for positive and negative feedback trials respectively, to model brain activity that co-varied with valence. The RPE regressor contained demeaned trial-by-trial signed RPEs from the Q-learning model. Since feedback valence and RPEs are correlated, i.e. positive feedback is accompanied by a positive RPE, this allowed us to assign brain activity co-varying specifically with valence or RPE. This was run as a fixed effects multiple regression model using FLAMEO on a per subject basis. Fixed effects multiple regression models for collapsing across runs and deriving within-patient medication difference COPEs were carried out in a similar way. Medication difference COPEs of feedback and RPE were then brought to the group level in a random effects model, using FSL’s FLAME 1+2 and outlier detection procedures. All group level Z (Gaussianized T) statistic images were thresholded using clusters of z > 2.3 and a cluster-corrected significance threshold of *p* < 0.01. Group-level analyses were carried out in this way for each regressor (feedback valence and RPE) on each separate group (HC, PD ON, PD OFF), on the within-subject medication differences ON > OFF and OFF > ON, and on across-subject disease differences HC vs PD (OFF or ON). All group-level z-statistic contrasts can be viewed at https://doi.org/10.6084/m9.figshare.6989024.v1. The MNI152_T1_1mm_brain standard brain was converted to functional space using FSL FLIRT, eroded by 1 voxel, and used as a brain mask for all of the analyses described above.

#### ROIs

We obtained high-resolution probabilistic atlas masks from a recent open-source dataset (Pauli et al., 2018). These sub-cortical ROIs have been well-established in playing an important role in reinforcement learning (Brown et al., 1999; Hazy et al., 2010; O’Doherty et al., 2017; Schultz et al., 1997). We focused on striatal ROIs (caudate nucleus, putamen, and nucleus accumbens) for the learning phase deconvolution analysis, as these have been most extensively studied in the past (Cools et al., 2007; Cox et al., 2015; Frank, 2005; Jahfari et al., 2017). FSL FLIRT was used to register the masks to FMRIPREP output space. BOLD percent signal change during the transfer phase was extracted from ROIs informed by the learning phase of the task. We took the cluster-corrected RPE medication difference (OFF-ON) z-statistic COPE from the learning phase and multiplied it by independent striatal ROIs from the Pauli et al. (2018) dataset. The bottom 25% of voxels from each of the resulting masks were excluded and the final masks were binarized. These masks are available at https://doi.org/10.6084/m9.figshare.6989024.v1.

#### Deconvolution analysis

Deconvolution analyses were carried out in striatal ROIs (see *ROIs* for mask information), to extract detailed BOLD time courses and tease apart the covariation with BOLD signal of positive and negative RPEs separately. We used the fMRI timeseries data already preprocessed by FMRIPREP. These timeseries were then converted to percent signal change (PSC). PSC was calculated by dividing the timeseries by the mean of the entire timeseries, multiplying by 100, and then subtracting 100, to get a mean-centered output timeseries. Data from each subject were weighted per voxel according to the probability of belonging to a particular striatal ROI, and then averaged across voxels of that ROI. We set up a model with three regressors: stimulus onsets (with RT duration), positive feedback onsets and negative feedback onsets. Positive and negative RPEs were z-scored separately and included as covariates of their respective positive or negative feedback event type. The deconvolution was implemented using the Python-based *nideconv* package (de Hollander and Knapen, 2017). Since we were not interested in the specific time courses related to stimulus onset, these events were modeled with a canonical HRF. Feedback events and covariates were deconvolved using a Fourier basis set, which uses a combination of sines and cosines to model the data. This was implemented instead of the standard finite impulse response (FIR) function as it substantially reduces the number of regressors, thereby improving the robustness of parameter estimates. Five Fourier regressors (1 intercept, 2 sine waves and 2 cosine waves) were used for each of the positive and negative feedback events and positive and negative RPE covariates. We also included several confound regressors in the model: FD, 6 rigid-body transform motion parameters (3 translational, 3 rotational), WM, stdDVARS (standardized derivative of RMS variance over voxels), and 6 aCompCor physiological noise regressors. Time courses were then estimated simultaneously using a least-squares fit, for the time window −2 to 13.05 seconds (7 TRs) around feedback onsets. Up-sampling during the fitting procedure was implemented 20-fold as part of *nideconv* functionality. The resulting time courses were then brought to the group level, and within-patient medication differences were calculated per up-sampled timepoint of each fit. Clusters of significant intervals were identified with *mne-python* (Gramfort et al., 2014, 2013) using permutation-based one-sample t-tests (t-threshold set at *p*<.05, n=5000 permutations). Shaded regions in Figure 4 and Figure S4 & S5 represent 68% confidence intervals (±1 SEM; bootstrapped using n=5000 permutations).

#### Transfer phase BOLD percent signal change

A standard GLM was set up to model BOLD responses to events in the transfer phase. Stimulus onsets and durations for three regressors were included: Approach A trials, Approach B trials, and all other trials (of no interest). Similar to the steps carried out in the learning phase whole-brain analysis, we performed 4mm smoothing, Savitsky-Golay high-pass filtering, and included the same 13 confound regressors in the design and convolved with a canonical HRF. Fixed effects analyses were performed across runs and for medication differences within patients. We then took the resulting two COPES for Approach A and Avoid B trials per subject and used FSL’s *featquery* to calculate the mean percent signal change in the striatal ROIs that showed a significant learning phase medication difference in RPE (described in *ROIs*). In a similar way to the behavioral correlation analysis between learning rate and transfer accuracy (*Behavioral analysis* and Figure 5B), we included learning rate EVs to explain the PSC (OFF>ON) medication difference in Avoid B>Approach A trials in striatal ROIs using robust (multiple) regression, with EVs as either: both positive and negative learning rate medication differences (kα_gain_ and kα_loss_), kα_gain_ only, or kα_loss_ only. Models were compared based on calculated BIC values (see Table S6), and the learning to transfer PSC correlation *p*-value was obtained from the winning α_loss_-only model. Individual medication differences were quantified as the modes of the within-subject medication difference parameter distributions.

## ACKNOWLEDGEMENTS

We would like to thank Annemarie Vlaar, Henk Berendse, and Odile van den Heuvel from the neurology and psychiatric departments of the Zaans MC and VUmc for help with patient recruitment, Ton Schweigmann and Joost Kuijer for practical and technical assistance at the MRI scanner, and Charlotte Koning, Daan Beverdam, and Rosa Broeders for help with data collection. We also thank Helen Steingroever and Ruud Wetzels for valuable insights on Bayesian modeling, Gilles de Hollander for inspiring discussions about fMRI modeling and Eduard Ort for exciting conversations about fMRI analyses and helpful comments on a version of the manuscript. This study was supported by an ERC Advanced Grant ERC-2012-AdG-323413 awarded to Jan Theeuwes.

## AUTHOR CONTRIBUTIONS

BM, SJ and TK designed the study. BM and GE collected the data. BM analyzed the data. SJ and TK supervised data analysis. BM, SJ and TK contributed (novel) methods. BM wrote the first draft of the manuscript. SJ and TK supervised and contributed to intermediate versions. BM, SJ, TK, GE and JT contributed to the final manuscript.

## CODE AVAILABILITY

Related analysis code scripts and output brain statistics are available to view at https://github.com/mccoyb4/Parkinson_RL.

## DATA AVAILABILITY

The data that support the findings of this study are available upon reasonable request from the corresponding author.

## SUPPLEMENTAL MATERIAL

**Figure S1.**
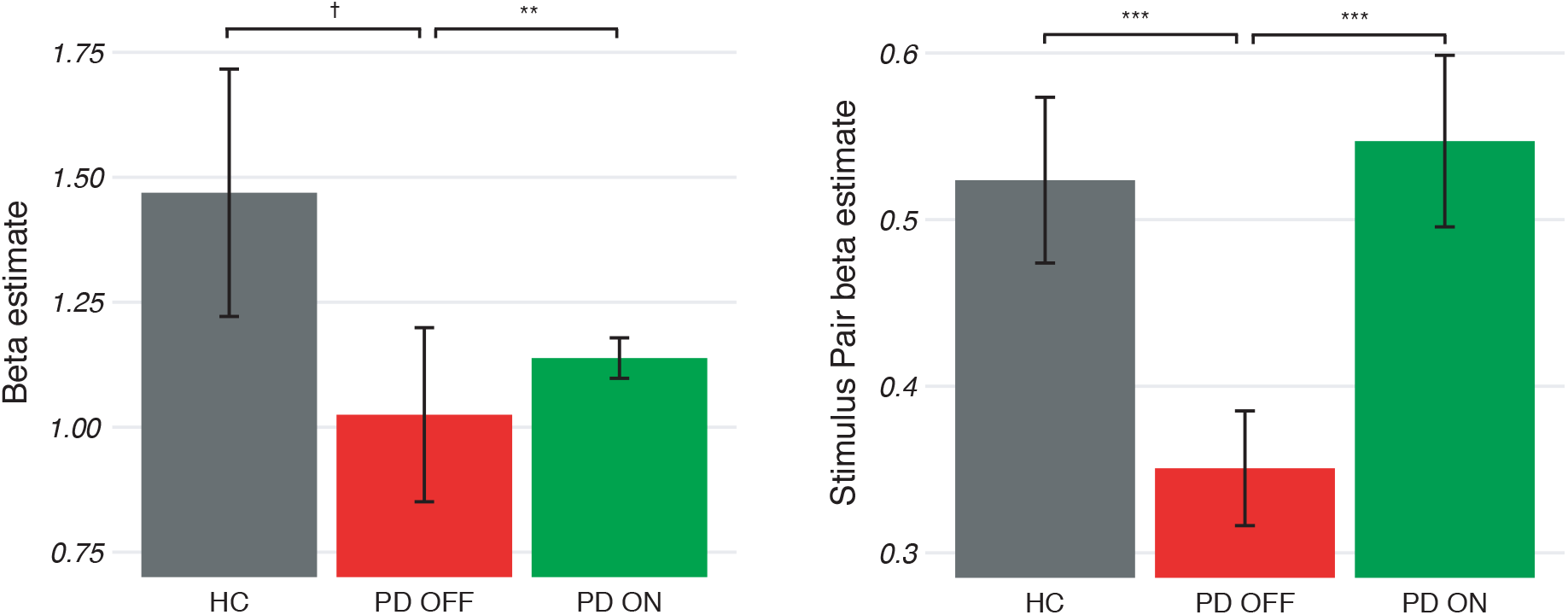
Beta parameter estimates for learning phase mixed-effects logistic regression model. PD OFF was considered as ‘baseline’, with any relative increase in beta parameters for PD ON or HC representing the effect of medication and disease status, respectively. Here, the main effects of disease and medication on choice accuracy are presented (left), as well as interaction effects of stimulus pair and disease, and stimulus pair and medication, on choice accuracy (right).

**Figure S2.**
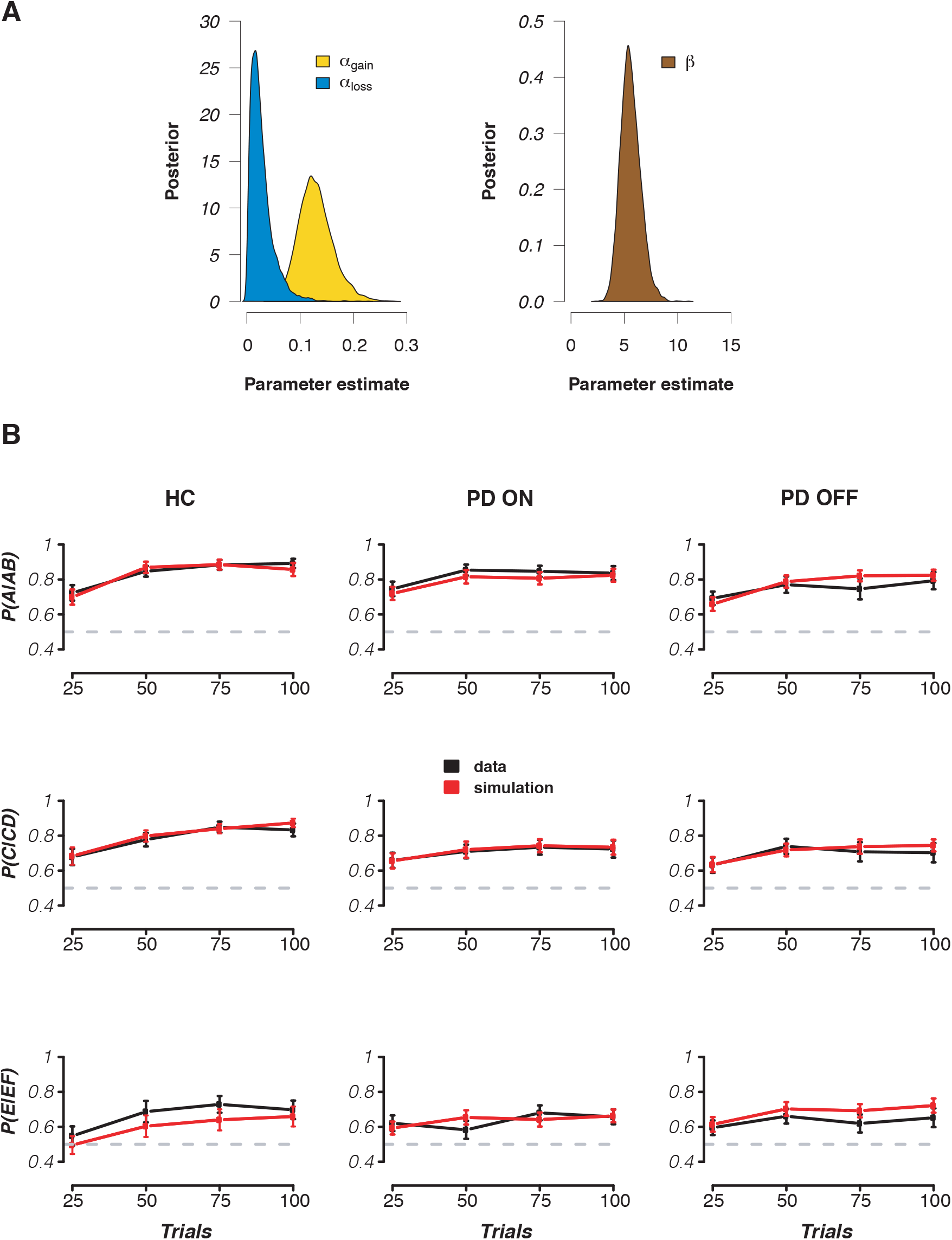
Bayesian hierarchical computational model assessment and simulation. **A)** Group-level parameter estimate distributions. **B)** Simulation of the fitted model. To check whether the model sufficiently captured actual choice behavior of participants, we simulated the probability of choosing the best option using the posterior distributions of the fitted free parameters of each participant. Plots of the modeled against empirical data across each group and stimulus pair show that the model is a good representation of overall learning.

**Figure S3.**
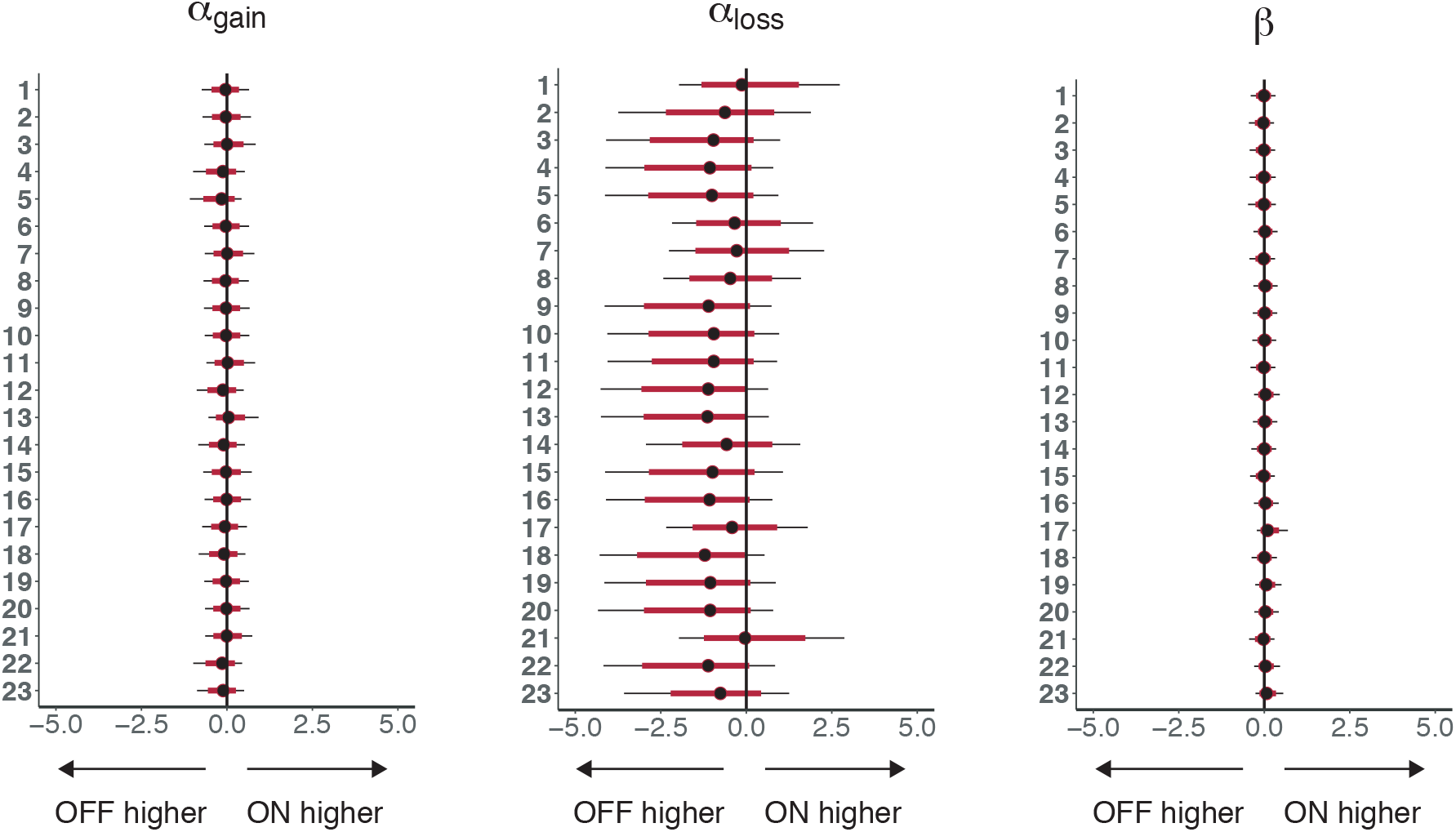
Individual medication differences in Parkinson’s patients in computational model parameters. Medians are displayed, with red and black horizontal bars denoting 85% and 95% highest density intervals (HDI), respectively. Note that wherever individual medication difference parameters are used, we take the mode rather than median of the distributions, as this point has the highest posterior density of all parameter values.

**Figure S4.**
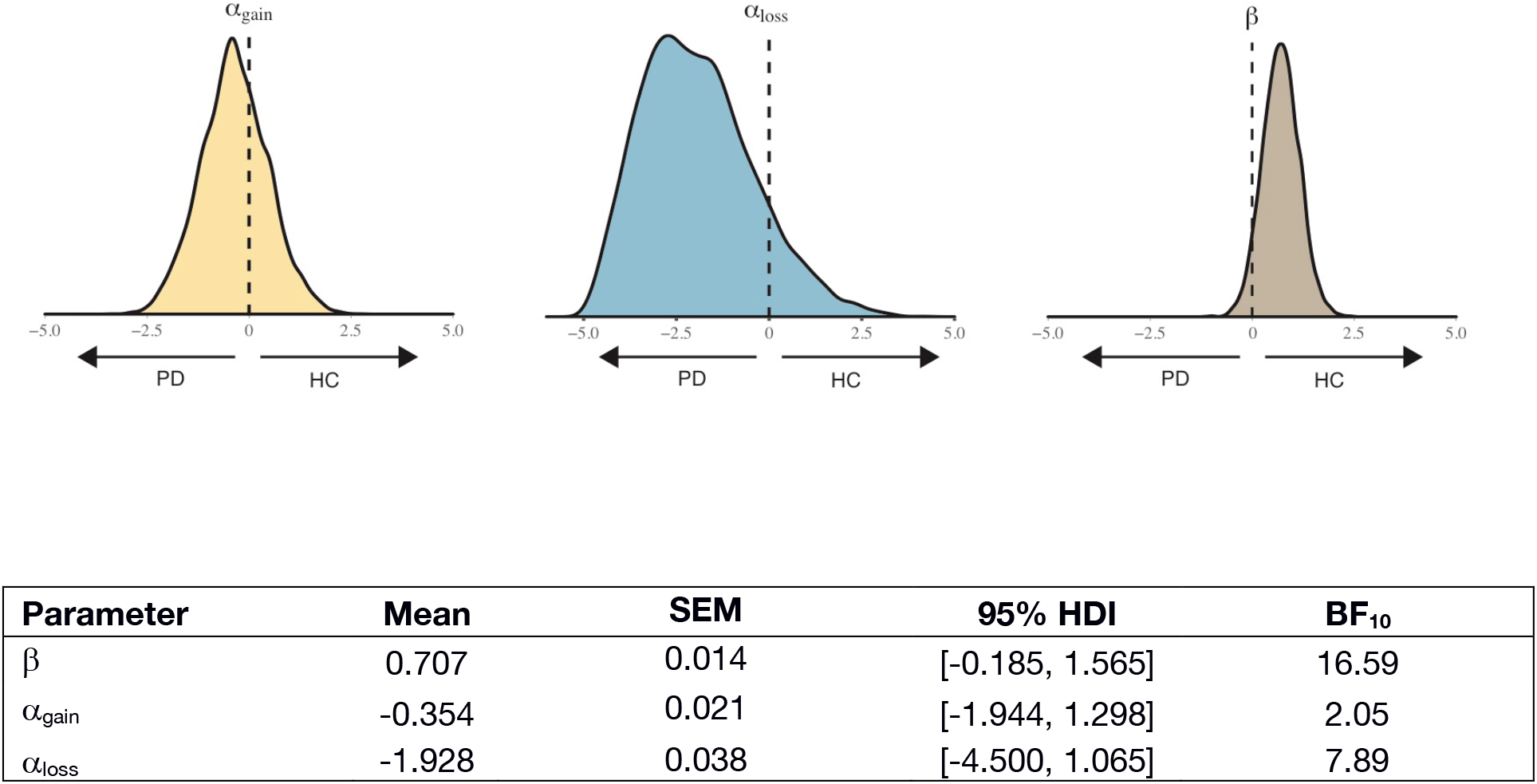
Disease differences in computational model parameters. Parameter distributions are shown for HC v PD (N=46). We found moderate evidence for greater α_loss_ negative learning in PD compared to HC (BF = 7.89), and strong evidence for greater exploitation in HC compared to PD patients (BF = 16.89).

**Figure S5.**
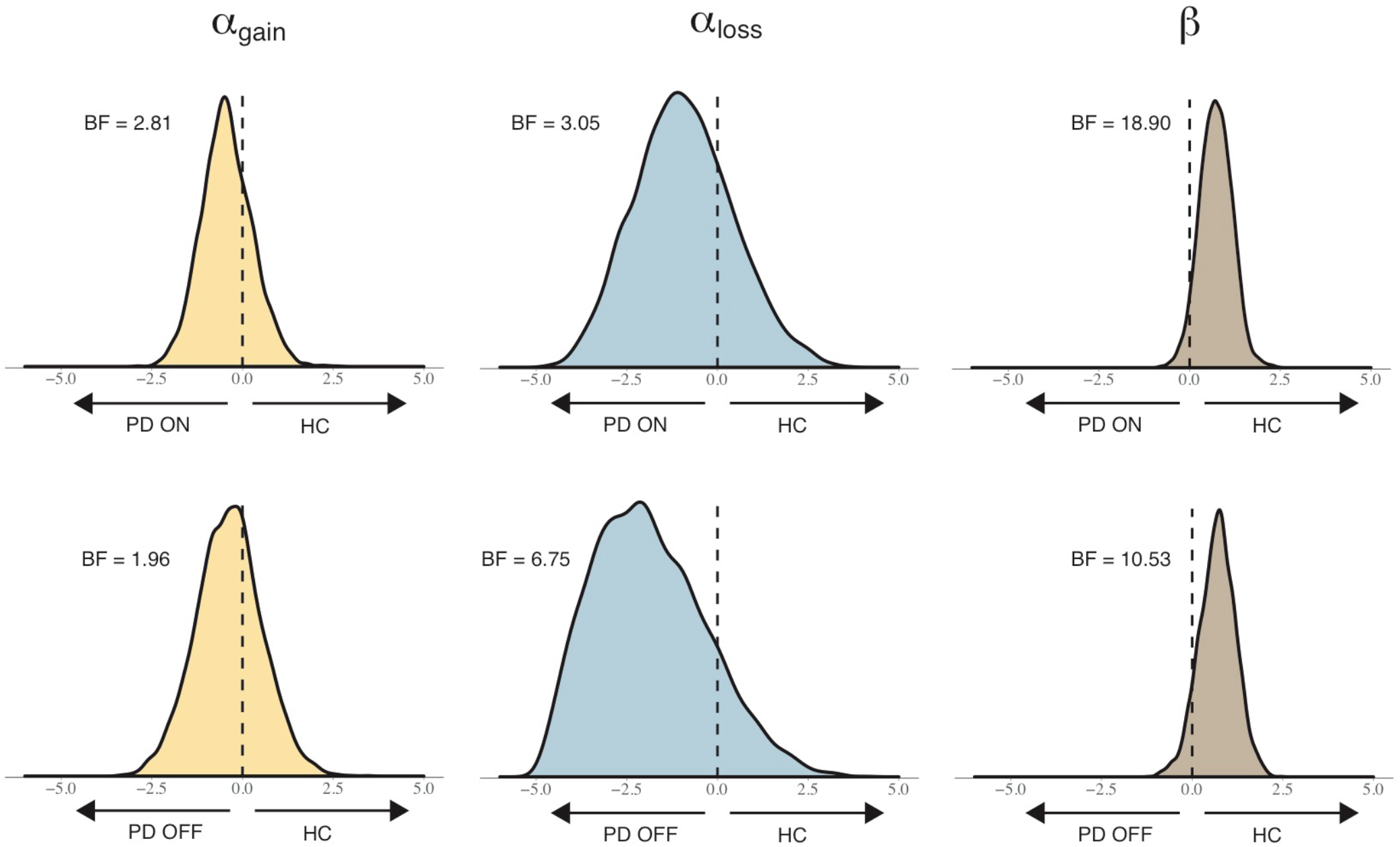
Disease differences in computational model parameters, separately for PD medication status. To assess whether both PD ON and OFF contributed in a similar or differential way to the disease-related difference distributions shown in Figure S4, two additional smaller models were run using a subset of data entered into the main model: HC vs. PD ON and HC vs. PD OFF. We ran 3000 samples each (discarding the first 1500 of each chain for burn-in). Checks for model convergence via visual inspection and *r̂* statistics was carried out in the same way as in the main model. The effect of disease on group-level parameters are displayed above for HC vs. PD ON (top row) and HC vs. PD OFF (bottom row). There is strong evidence for greater exploitation in HC compared to both PD ON and OFF, as indicated by a rightward shift in the β parameter distribution and BFs > 10.

**Figure S6.**
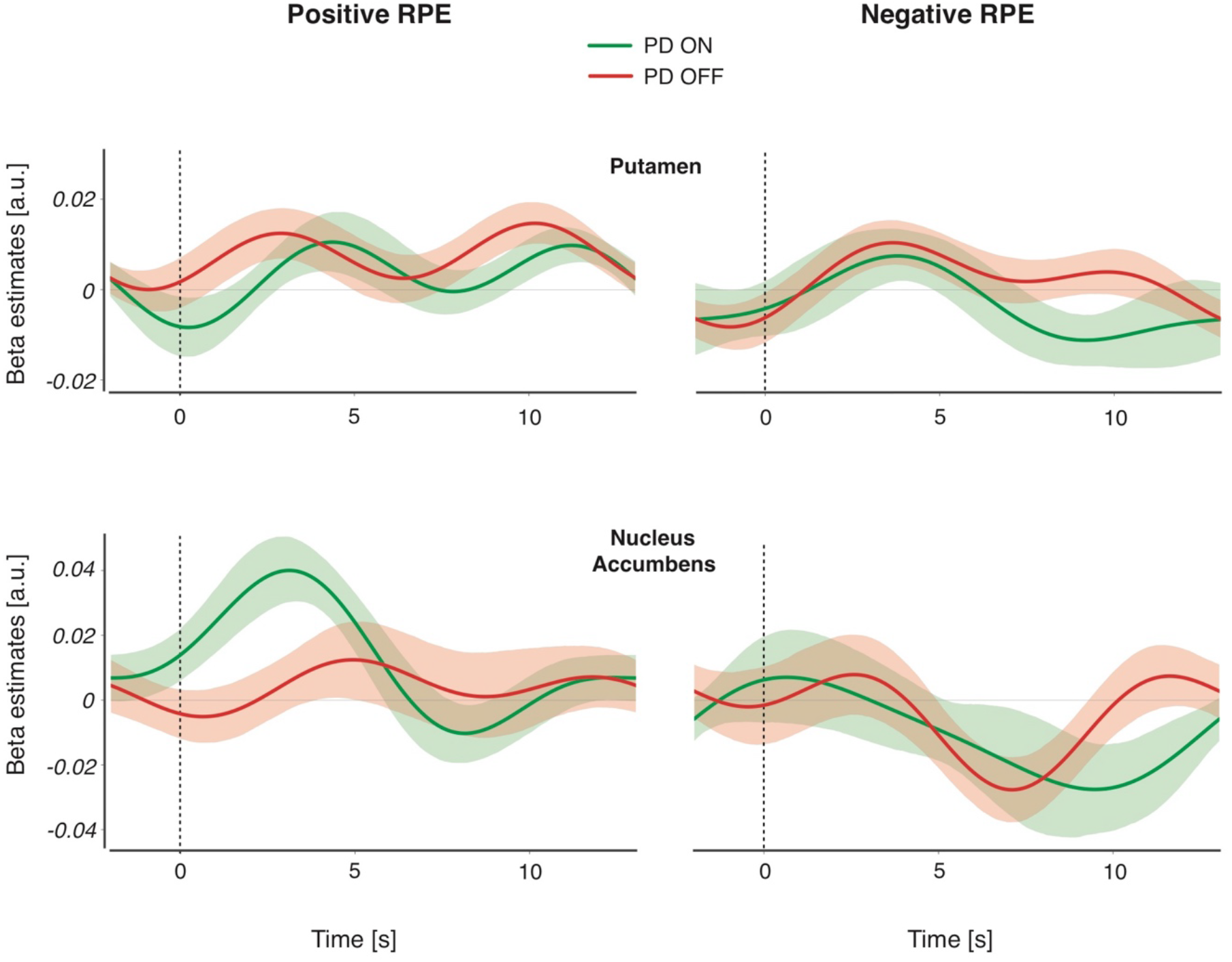
Positive and negative RPE in striatal ROIs. In addition to the caudate nucleus, we ran a deconvolution analysis in the putamen and nucleus accumbens (NAc). Positive and negative RPEs were z-scored around positive and negative events, respectively. Although no medication difference in RPE was found for ventral striatal NAc activity, it was included here for informational purposes since it has been implicated in several previous studies on the effects of dopamine on learning (Breiter et al., 2001; Cools et al., 2007; McClure et al., 2003; O’Doherty et al., 2003). In NAc, there appears to be a quantitative PD ON > OFF group level difference in positive RPE, however this difference is not statistically significant when within-subject differences were cluster-corrected across multiple timepoints.

**Figure S7.**
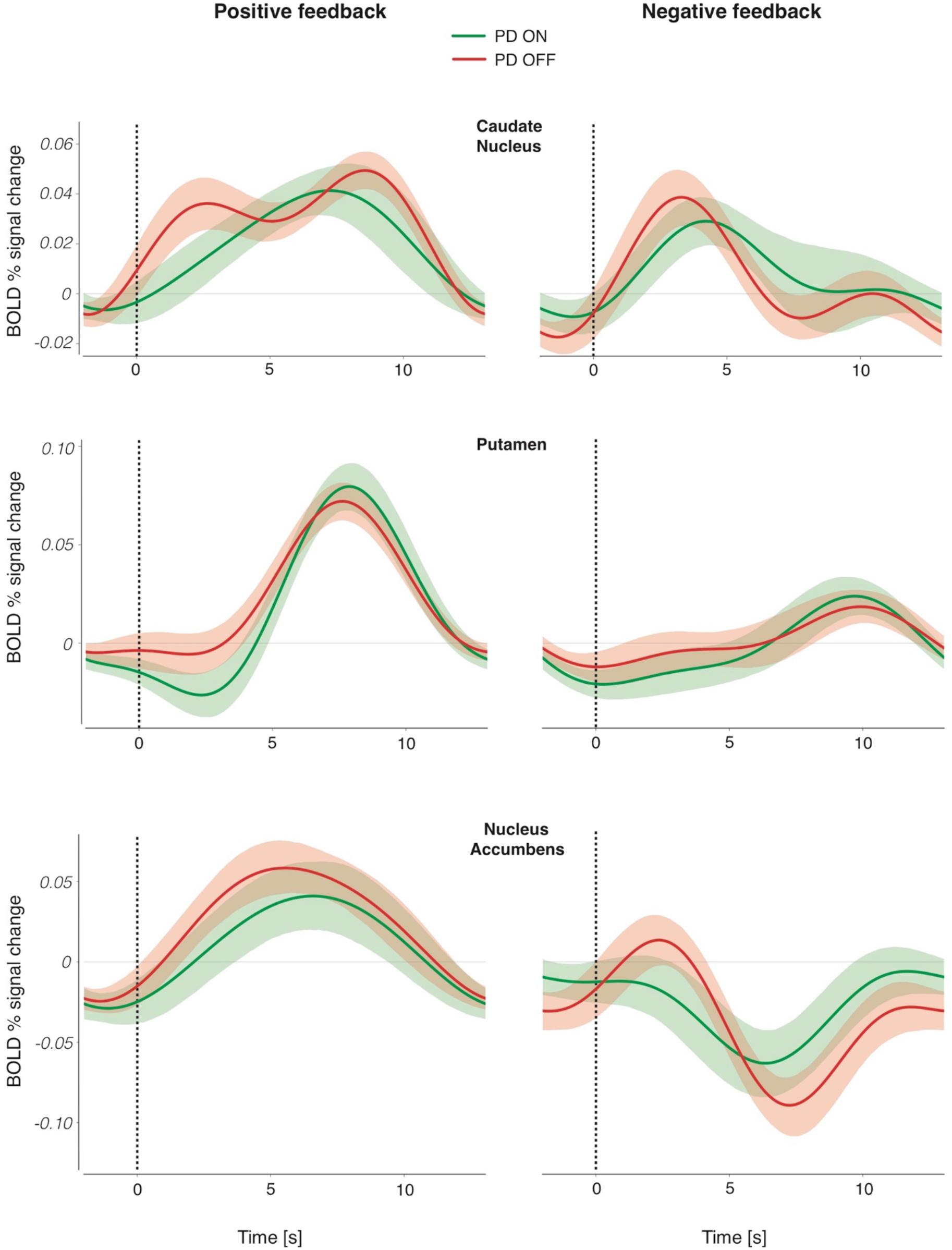
Changes in BOLD percent signal change for positive and outcomes in striatal ROIs. Percent signal change is shown for the caudate nucleus, putamen, and NAc. BOLD percent signal change is presented separately for positive and negative feedback events, for both PD ON (green) and PD OFF (red). Colored bands represent 68% confidence intervals.

**Figure S8.**
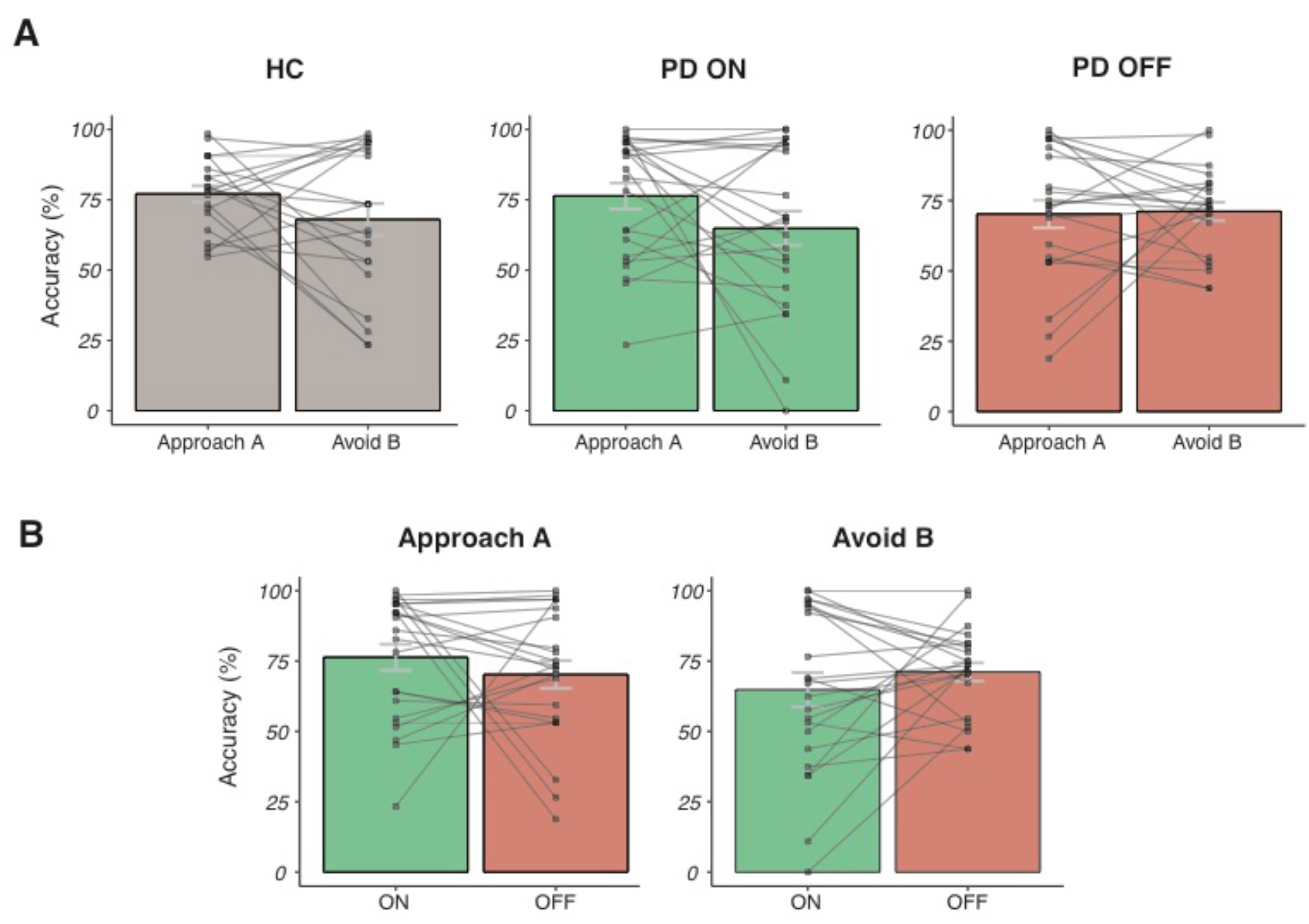
Subject-level accuracy plots in Approach A and Avoid B trials of the transfer phase. **A)** Averages per condition and within-subject plots, separately for HC, PD ON and PD OFF. **B)** Within-PD subject plots across medication session, separately per Approach/Avoid condition.

**Table S1.** Demographic and clinical characteristics of participants. Executive functioning was assessed using the following tests: the Montreal Cognitive Assessment (MoCA), the Dutch version (NLV) of the National Adult Reading Test (NART) as a measure of pre-morbid IQ, the Stroop color-word task to assess effects of interference, verbal (category) fluency tests, and the rule-shift cards test of the Behavioural Assessment of the Dysexecutive Syndrome (BADS), to assess mental flexibility (Wilson et al., 1997). The Complex Figure of Rey (CFR) was used as a measure of visuospatial memory. Verbal memory was assessed using the Dutch version of the Rey Auditory Verbal Learning Test (AVLT), testing both short- and long-term verbal memory (Saan and Deelman, 1986). Digit span forwards and backwards in short form (WAIS) was used to assess working memory. Participants also completed several self-report questionnaires: Beck Depression Inventory (BDI), Beck Anxiety Inventory (BAI), and Monetary Choice Questionnaire (MCQ). PD patients additionally completed the Wearing-Off Questionnaire (WOQ-Q10; related to the wearing off of DA medication), and the Questionnaire for Impulsive-Compulsive Disorders in Parkinson’s Disease-Rating Scale (QUIP-RS). The motor part of the Unified Parkinson’s Disease Rating Scale (UPDRS III) was carried out before each fMRI session. An overview of several test scores is provided in the table below. These assessments were not examined in the current study but are discussed in greater detail elsewhere (Engels et al., 2018; Engels et al., *in press*). Quantities are presented as the mean across the sample, with brackets denoting 1 standard deviation.

| Characteristic | Healthy Controls (HC)<br>N=23 | Parkinson patients (PD)<br>N=23 | HC vs. PD<br>(difference) |
| --- | --- | --- | --- |
| Age (years) | 60.35 (8.72) | 63.30 (8.24) | $t(44) = 1.182, p=.244$ |
| Gender | 8 females | 6 females | $\chi^2(1) = .411, p=.522$ |
| Education level (Verhage) | 6.09 (0.85) | 5.26 (1.14) | $t(44) = 2.793, p=.008^{**}$ |
| MoCA score | 27.91 (1.88) | 26.96 (1.92) | $t(44) = 1.708, p=.095$ |
| BDI | 4.09 (3.03) | 6.07 (4.31) | $t(44) = 1.802, p=.078$ |
| BAI | 23.17 (2.27) | 31.48 (6.15) | $t(44) = 6.077, p<.001^{***}$ |
| Digit Span Backwards | 7.00 (2.09) | 6.35 (2.39) | $t(44) = .986, p=.329$ |
| NLV reading score | 90.96 (8.47) | 89.57 (7.36) | $t(43) = .588, p=.560$ |
| Verbal fluency | 43.91 (11.26) | 37.91 (11.01) | $t(43) = 1.808, p=.078$ |
| Disease duration (years) | - | 4.00 (3.18) | - |
| UPDRS III on | - | 17.87 (7.77) | - |
| UPDRS III off | - | 20.65 (8.22) | - |
| Hoehn & Yahr | - | 2.11 (0.58) | - |
| LEDD (mg) | - | 790 (629) | - |

**Table S2.** Medication information for PD patients. Patient information regarding levodopa, dopamine agonists and any other dopaminergic medication. Levodopa Equivalent Daily Dosage (LEDD) was calculated according to Tomlinson et al., 2010.

| Patient (Nr.) | Age (years) | Disease duration (years) | LEDD (mg) | Medication information |  |  | Time to scan since medication (hours) |  |
| --- | --- | --- | --- | --- | --- | --- | --- | --- |
|  |  |  |  | Levodopa | DA-agonist | Other | OFF | ON |
| 1 | 55 | 3.5 | 564 | Yes |  |  | 15.0 | 1.0 |
| 2 | 73 | 2.0 | 752 | Yes |  |  | 17.0 | 1.0 |
| 3 | 67 | 10.0 | 564 | Yes |  |  | 20.5 | 1.0 |
| 4 | 72 | 3.0 | 375 | Yes |  |  | 16.5 | 1.5 |
| 5 | 68 | 1.0 | 828 | Yes |  |  | 16.0 | 2.5 |
| 6 | 56 | 2.0 | 375 | Yes |  |  | 26.5 | 2.0 |
| 7 | 68 | 2.0 | 378 | Yes |  |  | 19.5 | 5.5 |
| 8 | 65 | 8.0 | 850 | Yes |  | MAO-B inhibitor (Rasagiline)<br>COMT inhibitor (Entacapone) | 12.5 | 1.5 |
| 9 | 62 | 4.0 | 2780 | Yes |  | COMT inhibitor (Entacapone) | 14.5 | 3.0 |
| 10 | 64 | 5.5 | 982 | Yes | Pramipexol |  | 14.5 | 2.0 |
| 11 | 68 | 2.0 | 125 | Yes |  |  | 15.5 | 2.5 |
| 12 | 69 | 1.0 | 500 | Yes |  |  | 15.0 | 8.0 |
| 13 | 73 | 5.0 | 375 | Yes |  |  | 13.5 | 3.5 |
| 14 | 70 | 3.0 | 1548 | Yes |  | COMT inhibitor (Entacapone) | 15.5 | 1.5 |
| 15 | 71 | 6.0 | 1038 | Yes | Pramipexol |  | 8.5 | 5.5 |
| 16 | 47 | 6.0 | 1428 | Yes | Ropinirol |  | 14.0 | 1.0 |
| 17 | 48 | 0.5 | 1000 | Yes |  |  | 15.0 | 1.5 |
| 18 | 56 | 6.0 | 935 | Yes | Pramipexol | MAO-B inhibitor (Rasagiline) | 13.0 | 1.5 |
| 19 | 66 | 1.0 | 90 | Yes |  |  | 16.5 | 2.0 |
| 20 | 53 | 5.0 | 615 | Yes | Ropinirol |  | 19.5 | 1.5 |
| 21 | 72 | 13.0 | 1150 | Yes | Pramipexol | MAO-B inhibitor (Rasagiline) | 16.0 | 2.0 |
| 22 | 57 | 1.0 | 106 | No | Pramipexol |  | 18.5 | 4.5 |
| 23 | 51 | 6.0 | 1645 | Yes | Ropinirol | Amantadine | 14.0 | 2.5 |
| 24 | 61 | 1.5 | 108 | No | Pramipexol |  | 27.0 | 12.0 |

**Table S3.** Model comparison. We tested two hierarchical Bayesian models and used the model that best explained the data, as estimated by the lowest Bayesian information criterion (BIC) score on both a subject and whole model basis. As well as the free parameters α_gain_ α_loss_ and β included in the previously described model, we additionally included a perseverance (“stickiness”) parameter (π) in a separate model, to account for any bias in choosing the same stimulus of a pair regardless of the reward outcome (Kable and Glimcher, 2007; Schönberg et al., 2007). π was included in the softmax equation and was bounded as [-5, 5] according to previous research using this parameter (Wunderlich et al., 2012). The log-likelihoods were calculated per subject in the Stan model using the log categorical probability mass function. This was updated on a trial-by-trial basis to reflect whether a trial was correct given the probability of choosing that option. These log-likelihoods (LLH) were then extracted from the fitted model on a per subject basis and used to calculate the BIC according to:

| Parameters | BIC | AIC |
| --- | --- | --- |
| $\alpha_{\text{gain}}, \alpha_{\text{loss}}, \beta$ | 290.55 | 266.89 |
| $\alpha_{\text{gain}}, \alpha_{\text{loss}}, \beta, \pi$ | 565.11 | 533.57 |

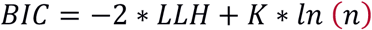

where K is the number of free parameters of the model and *n* is the number of observations (the number of trials, in this case). The BIC can also be calculated on a whole model basis, where LLH and *n* are the combined across all subjects. The table above shows the overall BIC values for each model; individual subject BICs are not shown here but are lower in all 46 subjects for the α_gain_, α_loss_, β model. AICs are also included in the table (AIC = −2 * *LLH* + 2*K*) to show that the α_gain_, α_loss_, β model is better in both cases.

**Table S4.** Summary of learning phase medication differences in posterior distributions of the Bayesian model group parameters. Bayes factors (BF) for medication differences in learning parameter distributions represent the BF according to direction of the visible shift in the posterior difference, i.e., the α_loss_ parameter in Figure 2B is shifted to the left (higher OFF medication), so the BF represents the probability of OFF > ON being greater than zero. HDI = highest density interval.

| Parameter | Mean | SEM | 95% HDI | BF<sub>10</sub> |
| --- | --- | --- | --- | --- |
| $\beta$ | 0.015 | 0.003 | [-0.213, 0.268] | 1.24 |
| $\alpha_{\text{gain}}$ | -0.048 | 0.005 | [-0.505, 0.407] | 1.39 |
| $\alpha_{\text{loss}}$ | -0.960 | 0.021 | [-2.566, 0.338] | 11.40 |

**Table S5.** Medication difference in whole brain RPE signal. Clusters of group-level PD OFF > ON medication difference (p <. 01, z = 2.3, cluster-corrected).

| # voxels | log10(p) | MAX X (mm) | MAX Y (mm) | MAX Z (mm) | COPE- MAX X (mm) | COPE- MAX Y (mm) | COPE- MAX Z (mm) | Areas included |
| --- | --- | --- | --- | --- | --- | --- | --- | --- |
| 529 | 25.6 | 12.3 | -82.8 | 47.4 | 35.2 | -72.9 | 54 | lateral occipital cortex, precuneus (R) |
| 233 | 13.7 | 5.71 | -89.3 | -15.3 | 2.43 | -79.5 | -8.7 | V1, occipital fusiform gyrus, occ. pole (R) |
| 101 | 6.75 | -33.7 | -43.4 | -45 | -30.4 | -43.4 | -45 | cerebellum (L) |
| 79 | 5.35 | -17.3 | -13.9 | 11.1 | -7.41 | 9.08 | 4.5 | caudate nucleus, putamen, glob. pallidus, thalamus (L) |
| 75 | 5.08 | -43.5 | -30.3 | 44.1 | -27.1 | -10.6 | 67.2 | postcent. gyrus, supramarg. gyrus, sup. parietal lobe (L) |
| 64 | 4.31 | 32 | -69.7 | -45 | -4.13 | -63.1 | -48.3 | cerebellum (R) |
| 44 | 2.78 | 51.6 | 35.3 | 30.9 | 38.5 | 45.2 | 37.5 | middle front. gyrus, frontal pole (R) |
| 42 | 2.62 | 61.5 | -43.4 | -8.7 | 61.5 | -46.7 | -8.7 | middle temp. gyrus, inf. temp. gyrus (R) |
| 42 | 2.62 | 28.7 | -7.32 | 50.7 | 32 | -4.04 | 50.7 | precent. gyrus, middle front. gyrus, sup. front. gyr (R) |
| 40 | 2.45 | 2.43 | 25.5 | 54 | 5.71 | 18.9 | 47.4 | sup. front. gyr. (R ) |
| 39 | 2.37 | 38.5 | -4.04 | 44.1 | 38.5 | 2.52 | 37.5 | precent. gyrus, middle front. gyrus, sup. front. gyr (R) |

**Table S6.** Summary of BIC values for the role of medication-related shifts in learning rate parameters in subsequent medication-related changes in transfer phase behavior and BOLD activity. BIC values relating to transfer phase behavior (second column) describe the explanatory power of within-patient medication-related shifts in learning rate parameters (kα_gain_, kα_loss_) in the transfer phase medication-related interaction in approach/avoidance behavioral accuracy. BIC values relating to brain activity (third column) describe the explanatory power of the same within-patient medication-related shifts in learning rate parameters in the transfer phase medication-related interaction in caudate nucleus BOLD activity during approach versus avoid trials. Overall, in both brain and behavior, the medication-related shift in only the negative learning rate, αloss, parameter best explained subsequent medication-related changes in approach/avoidance trials.

| Explanatory variable | BIC (behavior) | BIC (brain) |
| --- | --- | --- |
| $k\alpha_{\text{loss}}$ only | 231.38 | 33.36 |
| $k\alpha_{\text{gain}} + k\alpha_{\text{loss}}$ | 234.34 | 36.17 |
| $k\alpha_{\text{gain}}$ only | 234.48 | 37.92 |

